# Distinct monocyte replenishment kinetics of macrophages at the oral barrier restrict early-life bacterial colonisation

**DOI:** 10.64898/2026.11.23.767558

**Authors:** Kelly Wemyss, Rufus Daw, David Posner, Joshua Cox, Flora A. McClure, Siddharth Krishnan, Annie Whelan, Hayley M. Bridgeman, Tom McCarthy, Ian E. Prise, Zhaoyuan Liu, Florent Ginhoux, Rebecca Gentek, Tovah N. Shaw, John R. Grainger, Joanne E. Konkel

## Abstract

Resident tissue macrophages (RTMs) support tissue integrity and homeostasis. However, RTM in the gingiva, the mucosal barrier surrounding the teeth and the site of the prevalent inflammatory disease periodontitis, remain minimally explored. The gingiva experiences a unique combination of challenges, it is continuously exposed to both microbial stimuli and mastication-induced tissue damage; RTM could provide barrier reinforcement against both these challenges. Here we outline that no long-lived macrophage population could be identified within the gingiva. We demonstrate that gingiva macrophages are continuously replenished at a rate not impacted by the considerable microbial burden at this site. Despite microbe-induced signalling failing to determine macrophage replenishment, these signals did play key roles in re-shaping the macrophage network in early-life. We show that microbial-signals drive early-life increases in gingiva macrophages which are transient but vital to restrict the growth of the oral commensal community. Combined, our study delineates the biology of gingiva macrophages across life, outlining a developmentally restricted window during which these cells play key roles in regulating microbial colonisation.

**Significance Statement:** Resident tissue macrophages are key regulators of barrier homeostasis, yet their biology within the gingiva, a key oral barrier and site of the prevalent inflammatory disease periodontitis, remains poorly understood. We found that, unlike most tissues, the gingiva lacks a population of long-lived macrophages and instead relies on continuous macrophage replenishment. While microbial signals did not contribute to this macrophage replenishment rate, microbial cues initiated early-life remodelling of the gingiva macrophage network. We show that microbial-signals drive early-life increases in gingiva macrophages which are transient but vital to restrict oral microbial colonisation, revealing an unexpected developmental window in which macrophages shape host-microbiota interactions at a major mucosal barrier.

## Introduction

Our understanding of the mononuclear phagocyte system, and in particular their specialisation in barrier environments, has expanded considerably in recent years. It has become clear that macrophage ontogeny, turnover and functional capacity is exquisitely tailored to each barrier surface to effectively support barrier integrity. For example, in the gastrointestinal tract macrophages have been shown to mediate key protective roles including maintenance of tolerance to commensal microbes [1, 2], pathogen protection through bactericidal activity [3], reinforcing the epithelial cell barrier and mediating neuroimmune interactions [4, 5]. Despite this detailed knowledge for certain barrier sites, there is limited information on the development and activities of macrophages residing within the oral barrier, in particular the gingiva, the mucosal barrier surrounding the teeth.

Immune cells residing in the gingiva experience a unique combination of challenges during health. The gingiva is constantly exposed to dietary and airborne antigens, whilst hosting diverse commensal bacteria that must be effectively restrained within their niche. Yet, unlike other barrier sites, the gingiva also experiences barrier damage as a result of mastication [6]. Thus, immune control within the gingiva requires a concomitant activation of effective immunity and pro-resolution mechanisms; gingival macrophages could sit as central hubs co-ordinating both these activities. Understanding how macrophages contribute to gingiva health is important to ascertain, as loss of appropriate immune control at this site leads to development of periodontitis, the most common chronic inflammatory disease of humans, affecting about 12.5% of the global population [7].

Resident tissue macrophages (RTMs) can broadly be considered to be either generated during development or derived from circulating monocytes that are recruited from the blood to tissues, where they differentiate into macrophages [8–10]. Macrophages are heterogenous populations, exhibiting diverse dynamics (with some self-renewing and others being replenished by recruited monocytes over different time-scales [8, 11, 12]), functions and locations [13]. Recent work has outlined that the tissue niche into which monocyte-replenished macrophages dwell has a greater impact on macrophage function than origin [14], indicating a complex interplay of factors that continuously shape macrophage activity, longevity and contribution to effective immunity within different tissues. Again using the GI tract as an example, it was initially thought that macrophages in the GI tract were continuously and rapidly replenished by monocytes, recruited to the barrier due to the high microbial burden at this site [15]. However, we previously identified GI tract Tim4^+^ macrophages as a long-lived population, with low levels of monocyte replenishment [11, 16]. Consensus has now been reached that a long-lived macrophage population is resident at this site [17], yielding a macrophage network containing both short and long-lived populations and generated from both definitive and embryonic haematopoiesis, co-ordinately acting to support effective GI barrier function.

Although a number of studies have explored the ontogeny of Langerhans cells within a specific gingival niche – the epithelium [18, 19], to date the origin and long-term maintenance of macrophages resident within the gingiva has not been examined. Similarly, the functional capacities of macrophages within the gingiva are uncertain. During periodontitis macrophages in the gingiva have been shown to exhibit a pro-inflammatory phenotype, often described as M1 macrophages, and contribute to bone erosion [20, 21]. However, gingiva macrophages have also been reported to promote resolution of periodontal inflammation through the production of IL-10, TGFβ and DEL-1 [22–25]. In line with this, macrophage heterogeneity has been highlighted in single-cell RNA-sequencing approaches from both mouse and human gingiva [23, 26–29]. However, these studies largely focused on outlining alterations in total gingiva cells and cell-signalling during periodontitis, as opposed to comprehensively defining macrophage diversity, phenotype and function in either health or disease. Thus, despite concerted research effort to define RTMs at other sites, the identity and contribution of macrophages to gingiva health remain ill defined.

Here we sought to address this research gap, defining the origins and longevity of gingiva macrophages and probe their roles in barrier health. We outline a heterogenous macrophage population that turns over rapidly from monocytes. Indeed, in contrast to other barrier sites, we were unable to identify a long-lived population of macrophages in the gingiva. This rapid turnover was not driven by the high microbial burden present at this site, as no alterations in macrophage longevity were seen when microbial levels were reduced. We show that gingiva macrophages were subject to early-life cues, with a substantive increase in gingiva macrophages at 3 weeks after birth before rapidly reducing to adult levels by 4 weeks of age. This increase in gingiva macrophages was CCR2-dependent and driven in response to commensal colonisation. Macrophage heterogeneity was similar in the gingiva of adult and 3 week old mice but macrophages enriched in genes associated with responses to bacteria and bacterial-derived products were increased in the gingiva of 3 week old mice. In line with this, we demonstrate that this early-life recruitment of macrophages to the gingiva was vital to limit the biomass of the oral commensal community. Collectively we outline the developmental requirements of gingival macrophages and detail a unique macrophage network resident within this barrier that in adulthood is continuously replenished. We show this macrophage network is remodelled during development, with early-life CCR2-dependent macrophage expansion within the gingiva being vital to restrict the oral bacterial load. Our study provides new insights into the gingival barrier immune cell network, outlining macrophages as a key component of the host-microbe dialog in early life.

## Results

### Gingiva macrophages are short-lived but their turnover is not impacted by microbial signals

We characterised the macrophages within the gingiva of naïve adult mice, this included both the gingival epithelium and underlying connective tissue (lamina propria). We have previously shown that Ly6C^hi^ monocytes are resident within the gingiva at much higher proportions than macrophages, arising from diverse haematopoietic stem and progenitor cells (HSPC) that are resident in healthy gingiva [30]. Compared to the proportion of monocytes, a smaller population of macrophages was resident within the gingiva (**Fig. 1A**); this was true when gating macrophages using markers Ly6C, CD64 and MHC-II (**Fig. 1A**), as well as alternative strategies [31] (**Fig. S1A,B**). Macrophage identity was confirmed by examining cell morphology (**Fig. S1C,D**). Further flow cytometric phenotyping of the monocyte and macrophage populations demonstrated the macrophage population exclusively expressed F4/80, MerTK, CD64 and MHCII, all markers expressed by tissue macrophages, and higher expression of IBA-1 and CX3CR1 compared to the Ly6C^hi^ monocyte populations (**Fig. 1B**). These macrophages were distinct from the previously described Langerhans cells which reside in the gingiva epithelium [18], as they did not stain for Langerin or EpCAM (**Fig. S1E**). High-resolution confocal immunofluorescence imaging confirmed lower proportions of macrophages in the gingiva compared to monocytes (**Fig. 1C** and **Fig. S1F**).

**Figure 1:**
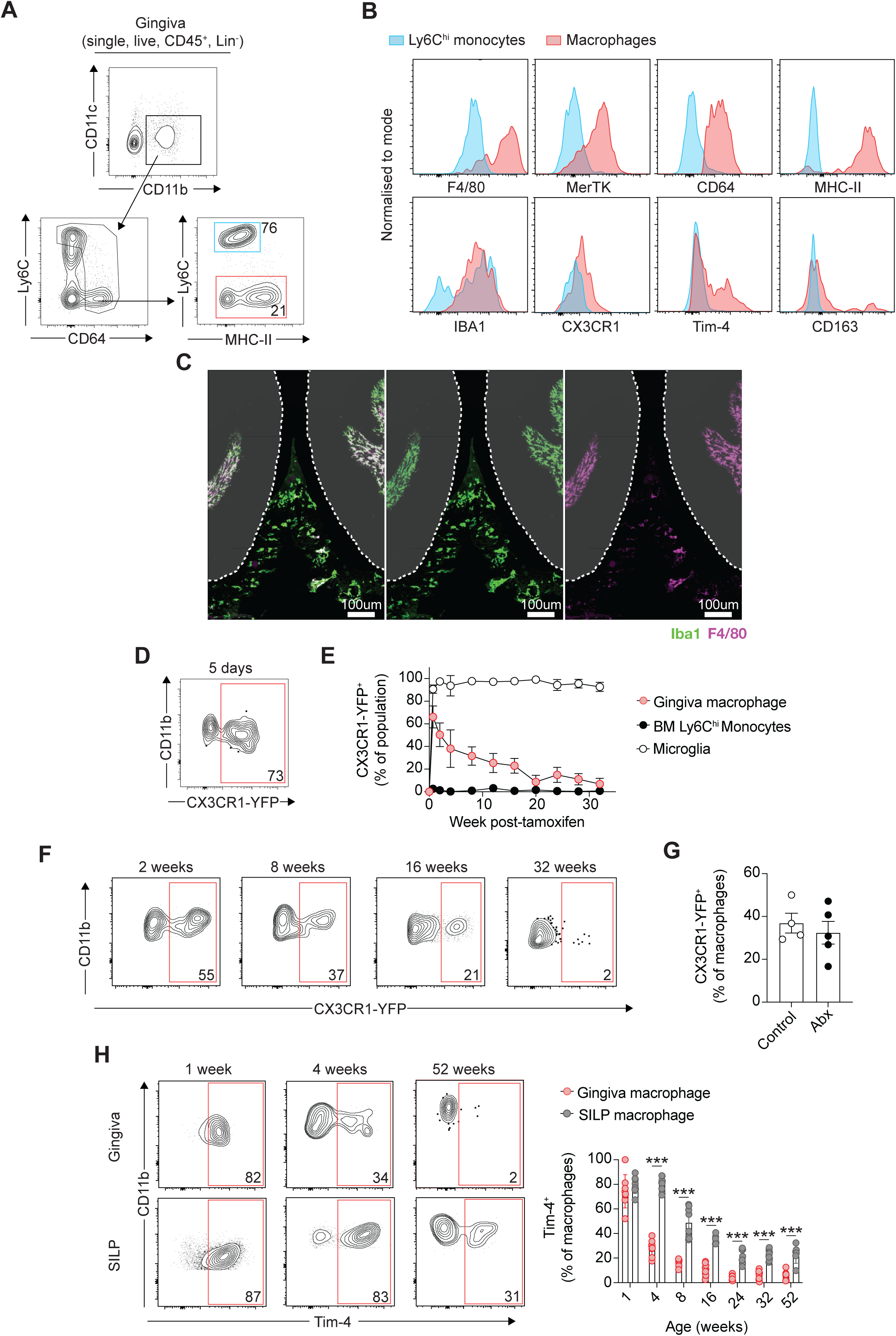
Gingiva macrophages are short-lived and their turnover is not impacted by local microbes. **(A)** Gating strategy to identify monocytes and macrophages in the gingiva. **(B)** Representative histograms showing staining for indicated markers on gingival monocytes (blue) and macrophages (red). **(C)** Confocal immunofluorescence microscopy images shown as maximum intensity projections of z-stacks. Iba1 (green) and F4/80 (purple) staining in the gingiva of adult C57BL/6 WT mice. Green cells indicate monocytes (Iba1^+^F4/80^-^), white cells indicate macrophages (Iba1^+^F4/80^+^). White dotted line indicates outline of each tooth. (**D**) *Cx3cr1^CreER^xR26-yfp* mice were treated with tamoxifen for 5 days, representative flow cytometry plot showing the frequency of YFP-expressing gingiva macrophages immediately after dosing. **(E)** Frequency of YFP-expressing gingiva macrophages, bone marrow (BM) Ly6C^hi^ monocytes and microglia at indicated time points post-tamoxifen treatment in *Cx3cr1^CreER^xR26-yfp* mice. **(F)** Representative flow cytometry plots showing the frequency of YFP-expressing gingiva macrophages in *Cx3cr1^CreER^xR26-yfp* mice 2 wk, 8 wk, 16 wk and 32 wk after tamoxifen treatment. **(G)** *Cx3cr1^CreER^xR26-yfp* mice were treated with antibiotics (Abx) for 4 wk following tamoxifen dosing. Frequencies of YFP-expressing gingiva macrophages in control and Abx treated animals. **(H)** Representative flow cytometry plots (left) and frequencies (right) indicating Tim-4 expression in gingiva and small intestine lamina propria (SILP) macrophages in C57BL/6 WT mice of indicated ages. Data are representative of at least two individual experiments, n=2-5 per experiment. Error bars show mean±SEM. Statistical comparisons were performed by Welch’s *t* test (D) or two-way ANOVA (F): *P≤0.05, ***P≤0.001.

Different organs and sub-tissular niches have distinct compositions of monocyte replenished macrophages and embryonically-derived RTM [13, 32–34]. Numerous studies and reviews have highlighted the gut as the organ with the largest proportion of macrophages rapidly replenished by bone marrow monocytes [11, 15, 17] and the brain as a site where embryonically-derived RTMs persist for a lifetime with only limited monocyte replenishment [8, 35]. Given the extremely high proportions of monocytes present in the healthy gingiva it is likely this key oral barrier is a site more akin to the gut in terms of monocyte-derived macrophage replenishment, but this has not been formally tested. To examine gingiva macrophage persistence and longevity we employed *Cx3cr1^CreER^xR26-yfp* mice in which administration of tamoxifen irreversibly induces eYFP expression by CX3CR1^+^ cells. Adult mice were treated with 5mg tamoxifen daily by oral gavage for 5 days and tissues were collected at defined timepoints following the final dose. Any cells developing from monocytes following the withdrawal of tamoxifen will be eYFP^-^, whilst eYFP^+^ cells will have persisted since tamoxifen administration. We observed substantial labelling of gingiva macrophages at 5 days post-tamoxifen (**Fig. 1D**) but, as expected, minimal labelling of gingival Ly6C^hi^ monocytes (**Fig. S2A**) [10]. As a control we saw that at all time points following labelling CX3CR1^+^ microglia persisted as eYFP^+^ cells (**Fig. 1E**). Although well labelled after treatment, gingival macrophages had become largely eYFP^-^by 20 weeks post labelling, gradually losing the eYFP-positive population (**Fig. 1E,F**). The half-life of eYFP^+^ cells was 75 days and, as such, so few persisted at later time points that specific marker expression was difficult to examine (**Fig. 1F**). These data suggest that macrophage turnover in the gingiva was fairly rapid and that, unlike almost every other tissue, there was no long-lived population of macrophages persisting within the gingiva. A caveat to this statement is that only about 75% of gingival macrophages label following tamoxifen treatment. However, this approach has been leveraged to examine macrophage longevity across barriers and organs, detailing a long-lived population in each case [10, 11].

Signals derived from local commensal microbial communities have been shown to contribute to macrophage turnover at other barrier sites [15], therefore we queried whether the oral bacterial community contributed to driving the continued gingiva macrophage replenishment. Utilising the *Cx3cr1^CreER^xR26-yfp* mice with the same tamoxifen dosing as above, we administered broad-spectrum antibiotics (ABX) for 4 weeks following tamoxifen treatment and before examining macrophage eYFP-labelling. This ABX-treatment protocol led to a substantive reduction in oral bacterial load (**Fig. S2B**). Importantly, reduction in oral bacterial burden had no impact on the loss of eYFP^+^ labelling within gingival macrophages (**Fig. 1G**), indicating that the bacterial burden in the gingiva does not drive the elevated replenishment rates seen.

Finally, we examined gingiva macrophage expression of Tim4, which has been shown to mark populations of long-lived macrophages across the body [11, 36, 37], including liver Kupffer cells, large peritoneal macrophages and long-lived gut macrophages. We examined proportions of Tim4^+^ macrophages in the gingiva from neonatal to aged time points. Similar to the small intestine, at early time points after birth a large majority of gingival macrophages were Tim4^+^ (**Fig. 1H**). However, this population was rapidly lost and by 8 weeks of age less than 20% of gingival macrophages were Tim4^+^ (**Fig. 1H**), with the majority of the loss occurring by 4 weeks of age. This contrasted the slow loss of Tim4^+^ macrophages within the small intestine and again suggests elevated replenishment rates of macrophages in the gingiva. Collectively our data suggest that macrophages residing within the gingiva possess distinct dynamics compared to other barriers, in that we could not identify a long-lived population and these macrophages were subject to continuous replenishment at a rate not shaped by microbial-derived signals.

### Macrophages in the gingiva are derived from definitive haematopoiesis

To further expand our understanding of the macrophage network in the gingiva we examined their ontogeny leveraging *Cdh5-*CreERT2.Rosa26^tdT^ mice, which enable fate mapping of the progeny of both yolk sac- and definitive haematopoiesis-derived cells in a precise manner (**Fig. 2A**) [38]. Mice were administered a single dose of 4-Hydroxytamoxifen (4-OHT) at E7.5 to label yolk sac derived cells and E10.5 to label the progeny of definitive haematopoiesis. Frequency of labelling was assessed in adult (8-12 week) and 1 year old mice. In mice labelled at E7.5 we saw no labelling of either monocytes or macrophages in the gingiva, indicating that neither population originates from yolk sac progenitors (**Fig. 2B**). This is in contrast to brain microglia which were well labelled in the offspring of mothers receiving 4-OHT at E7.5 (**Fig S2C**). In adult mice labelled at E10.5, we observed that all gingiva monocytes were labelled (**Fig. 2C**), as were blood monocytes (**Fig. S2C**). In these same animals approximately 60% of gingival macrophages displayed labelling, indicating that adult gingiva macrophages are partially derived from adult haematopoietic stem cells (HSC) and are partially of embryonic origin. Given the lack of labelling in these cells at E7.5, gingiva macrophages could potentially be partially of foetal liver origin. By 1 year of age, macrophages also show an equivalent level of labelling as gingiva and blood monocytes, indicating replacement of embryonic derived cells with cells derived from definitive haematopoiesis (**Fig. 2C**).

**Figure 2:**
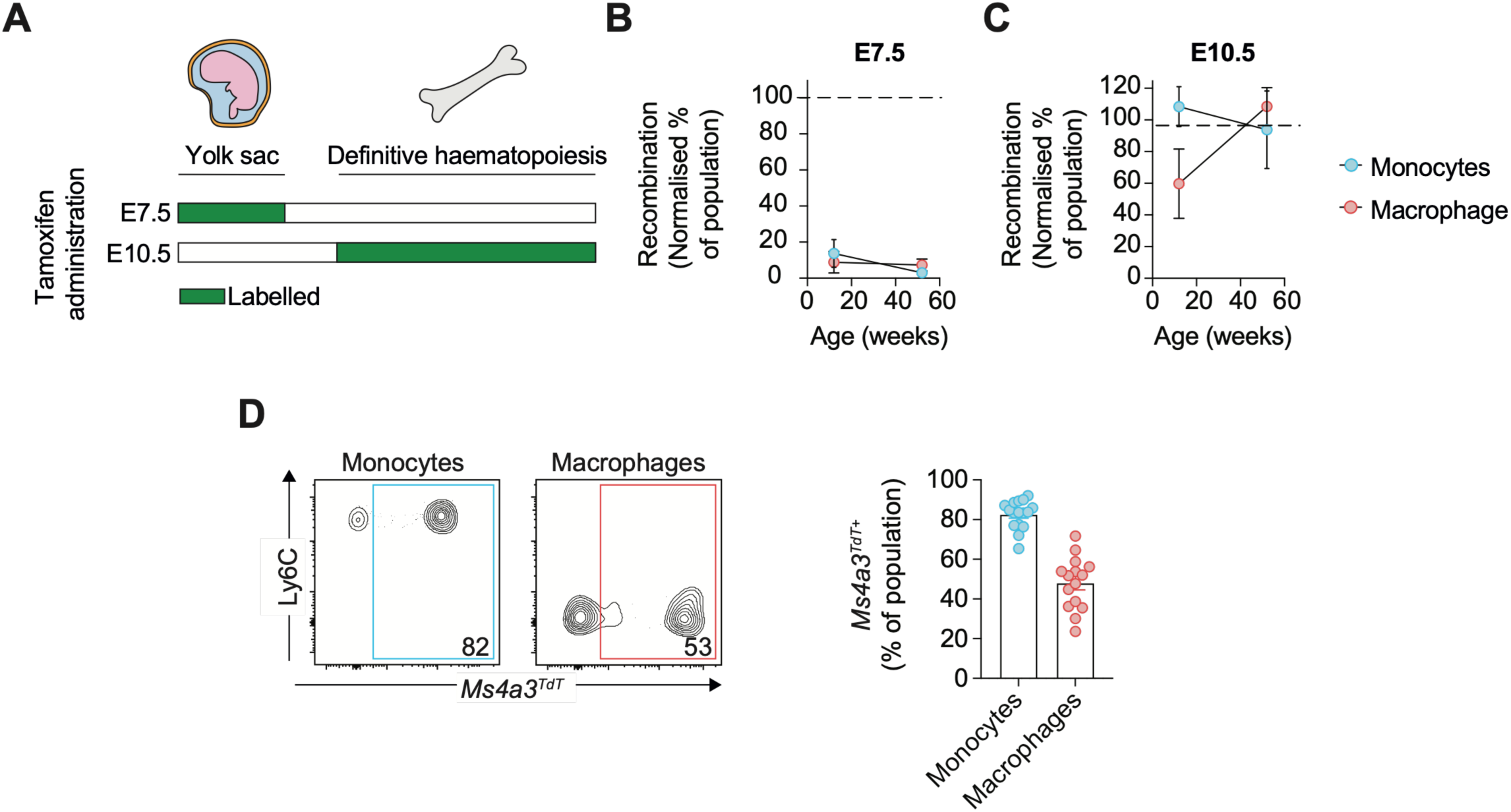
Gingiva macrophages are derived from definitive haematopoiesis. **(A)** Schematic showing labelling of cells of yolk sac or definitive haematopoiesis origin following tamoxifen treatment at embryonic day 7.5 (E7.5) or 10.5 (E10.5) in *Cdh5*-CreERT2:Rosa^tdT^ mice. **(B,C)** Frequencies of reporter recombination by gingival monocytes and macrophages in *Cdh5*-CreERT2:Rosa^tdT^ mice pulsed with 4OHT at **(B)** E7.5 or **(C)** E10.5. **(D)** Representative flow cytometry plots (left) and frequencies (right) indicating expression of *Ms4a3^TdT^*in gingiva monocytes and macrophages in 8-wk-old *Ms4a3^TdT^* reporter mice. Data are representative of at least two individual experiments, n=2-5 per experiment. Error bars show mean±SEM.

Complimentary to this, we also examined gingiva macrophages in *Ms4a3^Cre^-Rosa^tdt^* reporter mice which label monocyte-derived cells [12]. In adult reporter mice at 8-12 weeks of age, ∼50% of gingiva monocytes were tdTomato-positive (**Fig. 2D**). These data outline that, similar to the Cdh5-labelling, in adult gingiva over half of all macrophages were derived from recruitment of conventional, Granulocyte-monocyte progenitor (GMP)-derived monocytes.

### Developmentally restricted macrophage expansion

Our previous work has shown a slow and gradual loss of Tim4^+^ macrophages in the gastrointestinal tract as recruited monocyte-derived cells progressively replenished tissue macrophage niches [11, 16]. However, this was not seen in the gingiva, where we saw a rapid loss of Tim4^+^ macrophages over the neonatal period (**Fig. 1H**). Shifts in the dynamics of monocyte replenishment of tissue macrophages have been previously described during early life as tissues develop [39, 40]. Our data showing substantive loss of Tim4^+^ gingiva macrophages during early life could indicate there is some dynamic remodelling in the gingiva macrophage network over this period. Indeed, there are a number of changes occurring at the gingiva within the first month of life; tooth eruption occurs between 2 and 3 weeks of age [41], resulting in tissue damage that requires rapid repair. This occurs on a background of commensal colonisation, and subsequent bacterial diversification, that is occurring after birth [42]. To examine this neonatal window further we examined macrophages in the gingiva over this period. We first enumerated macrophages and monocytes across the neonatal period and into adulthood. We observed an overall increase in macrophage numbers at 3 weeks, which was followed by a fall in numbers by 4 weeks of age to a level which they then persisted into adulthood (**Fig. 3A**). This macrophage increase was slightly preceded by a monocyte increase peaking at 16 days post birth (**Fig. 3B**). The increase in gingiva macrophages at 3 weeks of age was due to an increase in MHCII^-^macrophages (**Fig. 3C,D**), with MHCII^+^ macrophages remaining at a consistent level post birth. This transient increase in macrophages was confirmed through high-resolution confocal immunofluorescence imaging, in which more macrophages can been seen within the gingiva of 3 week old mice (**Fig. 3E** and **Fig S3A**) compared to adults (**Fig. 1C**).

**Figure 3:**
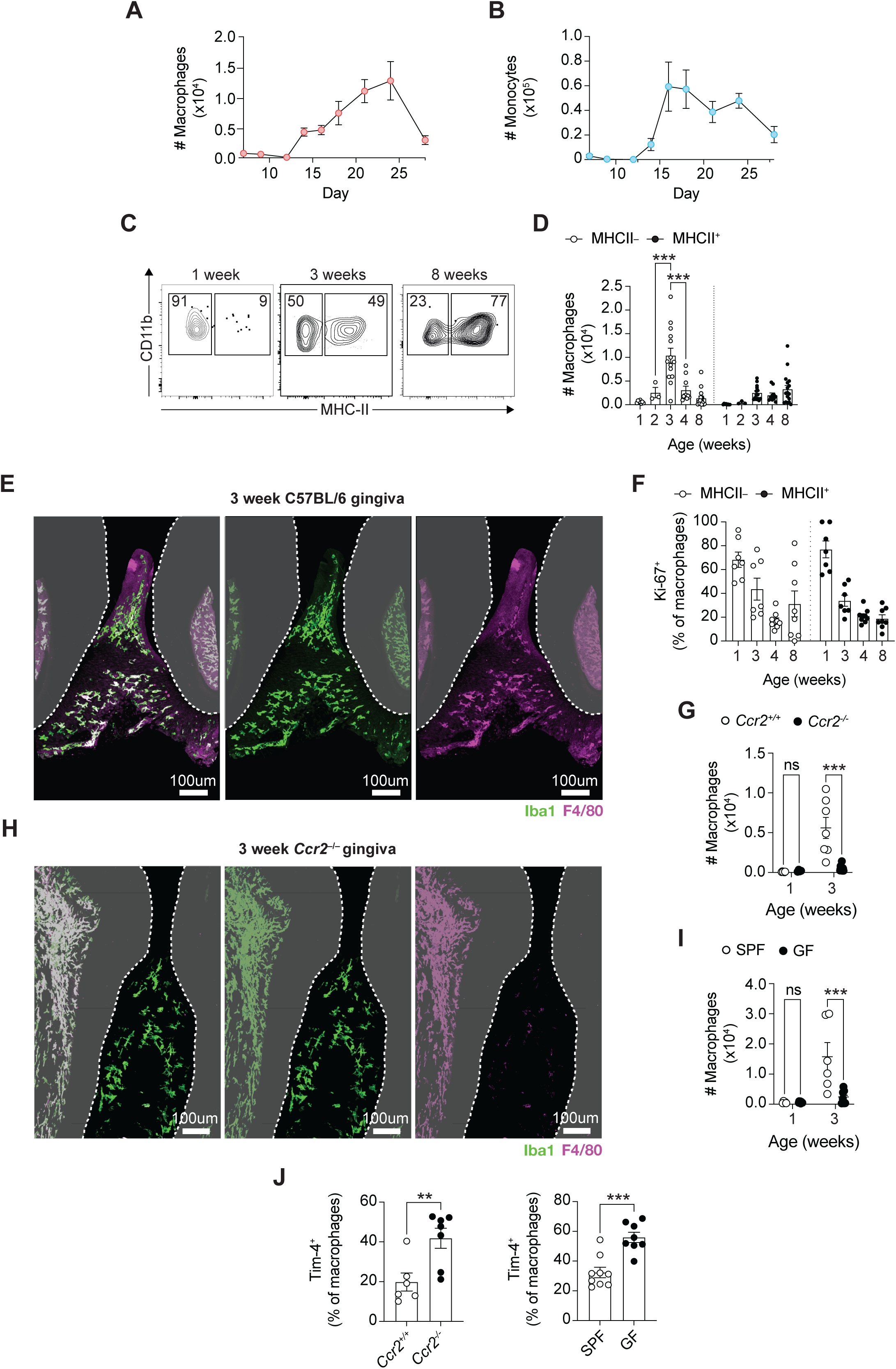
Microbial stimuli drives transient increases in gingiva macrophages over the neonatal period. **(A,B)** Absolute numbers of **(A)** gingival macrophages and **(B)** gingiva monocytes in C57BL/6 WT mice of indicated ages. **(C)** Representative flow cytometry pots indicating expression of CD11b and MHC-II in gingival macrophages in C57BL/6 WT mice of indicated ages. **(D)** Absolute numbers of MHC-II^-^and MHC-II^+^ gingival macrophages in C57BL/6 WT mice of indicated ages. **(E)** Confocal immunofluorescence microscopy images shown as maximum intensity projections of z-stacks. Iba1 (green) and F4/80 (purple) staining in the gingiva of 3-week old C57BL/6 WT mice. Green cells indicate monocytes (Iba1^+^F4/80^-^), white cells indicate macrophages (Iba1^+^F4/80^+^). White dotted line indicates outline of each tooth. **(F)** Frequency of Ki-67 expressing gingival MHC-II^-^ and MHC-II^+^ macrophages in C57BL/6 WT mice of indicated ages. **(G)** Absolute numbers of gingival macrophages in 1 and 3-wk old *Ccr2^+/+^* and *Ccr2^−/−^* mice. **(H)** Confocal immunofluorescence microscopy images shown as maximum intensity projections of z-stacks. Iba1 (green) and F4/80 (purple) staining in the gingiva of 3-week old *Ccr2^−/−^* mice. Green cells indicate monocytes (Iba1^+^F4/80^-^), white cells indicate macrophages (Iba1^+^F4/80^+^). White dotted line indicates outline of each tooth. **(I)** Absolute numbers of gingival macrophages in 1 and 3-wk old SPF and GF mice. **(J)** Frequency of Tim-4^+^ gingiva macrophages in 3-wk-old *Ccr2*^+/+^ and *Ccr2^−/−^* mice (left) and 3-wk-old SPF and GF mice (right). Data are representative of at least two individual experiments, n=2-5 per experiment. Error bars show mean±SEM. Statistical comparisons were performed by two-way ANOVA (D, F, G, I) or Welch’s *t* test (J): **P≤0.01, ***P≤0.001.

We next examined the driver of macrophage expansion at 3 weeks of age. We first examined macrophage proliferation, staining for Ki67. At early life timepoints more macrophages in the gingiva were Ki67^+^, with proportions of Ki67^+^ macrophages dropping at 4 weeks of age and persisting at those levels into adulthood (**Fig. 3F**). However, proportions of Ki67^+^ macrophages were similar in MHCII^+^ and MHCII^-^ gingiva macrophages, indicating that the MHCII^-^macrophage expansion was not exclusively due to proliferation. We next examined whether monocyte recruitment could be contributing to the gingiva macrophage increase at 3 weeks. To explore this we examined the macrophage population in 3 week old *Ccr2^−/−^*mice and littermate controls. The macrophage increase seen at 3 weeks of age was completely abolished in *Ccr2^−/−^* mice, seen via both flow cytometry and imaging approaches (**Fig. 3G,H** and **Fig. S3B**). Thus, the expansion in MHC-II^-^macrophages within the gingiva of neonatal mice was driven by CCR2-mediated monocyte recruitment into the tissue.

Next, we considered the upstream signal driving CCR2-mediated monocyte recruitment to the gingiva at 3 weeks. To investigate the role of the microbiome as the stimulus for monocyte recruitment we examined macrophages numbers in germ free (GF) mice. We observed no increase in macrophage numbers at 3 weeks of age in GF animals (**Fig. 3I**). These data suggest a microbiome driven increase in monocyte-derived macrophages in the gingiva in early life. This increase is transient as both *Ccr2^−/−^*and GF mice exhibited similar numbers of gingiva macrophage to controls by adulthood (**Fig. S3C**). In line with this transient, early-life increase in monocyte-derived macrophages, a higher proportion of gingiva macrophages present at 3 weeks in both *Ccr2^−/−^* and Germ Free mice were Tim4^+^ (**Fig. 3J**), indicating the early life loss of Tim4^+^ macrophages is driven by microbiome driven monocyte-replenishment of RTM in the gingiva.

### Macrophages in the gingiva of 3 week old mice are enriched in bacterial response genes

Our data indicate an acute expansion in MHC-II^-^macrophages resident in the gingiva barrier at 3 weeks of age, before this population contracts again by 4 weeks, continuing at this size into adulthood. To further characterise the expanded macrophage population compared to that in adult gingiva we performed single-cell RNA-sequencing of macrophages purified from the gingiva of naive 3 week old and adult mice. Although gingiva macrophages could be subdivided into 8 clusters (**Fig. 4A,B**), with all clusters expressing macrophage defining lineage markers *Adgre1, C1qa, Csfr1* and *Cx3cr1* (**Fig. S4A**), they separated into two distinct populations (Clusters 1 and 6 forming population A and Clusters 0, 2, 3, 4, 5, 7 forming population B). Broadly, population A was enriched in *Malat1*, *Mef2a*, *Atp2b1* and *Fyb* (**Fig. 4C**), all genes associated with proinflammatory macrophage phenotypes [43–45]. Yet within this population, cluster 6 was clearly distinct, enriched in *Ccr2, Nfkb1*, *Irf4*, *Nr4a1*, and *Cebpb*, genes increased in monocytes suggesting recent recruitment and development of this population from monocytes (**Fig. S4B**). Population B was not only more diverse but contained more cells. Pseudotime analysis on population B revealed a branched trajectory progressing from cluster 3, a population enriched in genes associated with the cell cycling including *Cdk1*, *Ccna2, Ccnb2,* and *Pcna (***Fig. S4B**), through cluster 2 and then cluster 0 (**Fig. S4C**).

**Figure 4:**
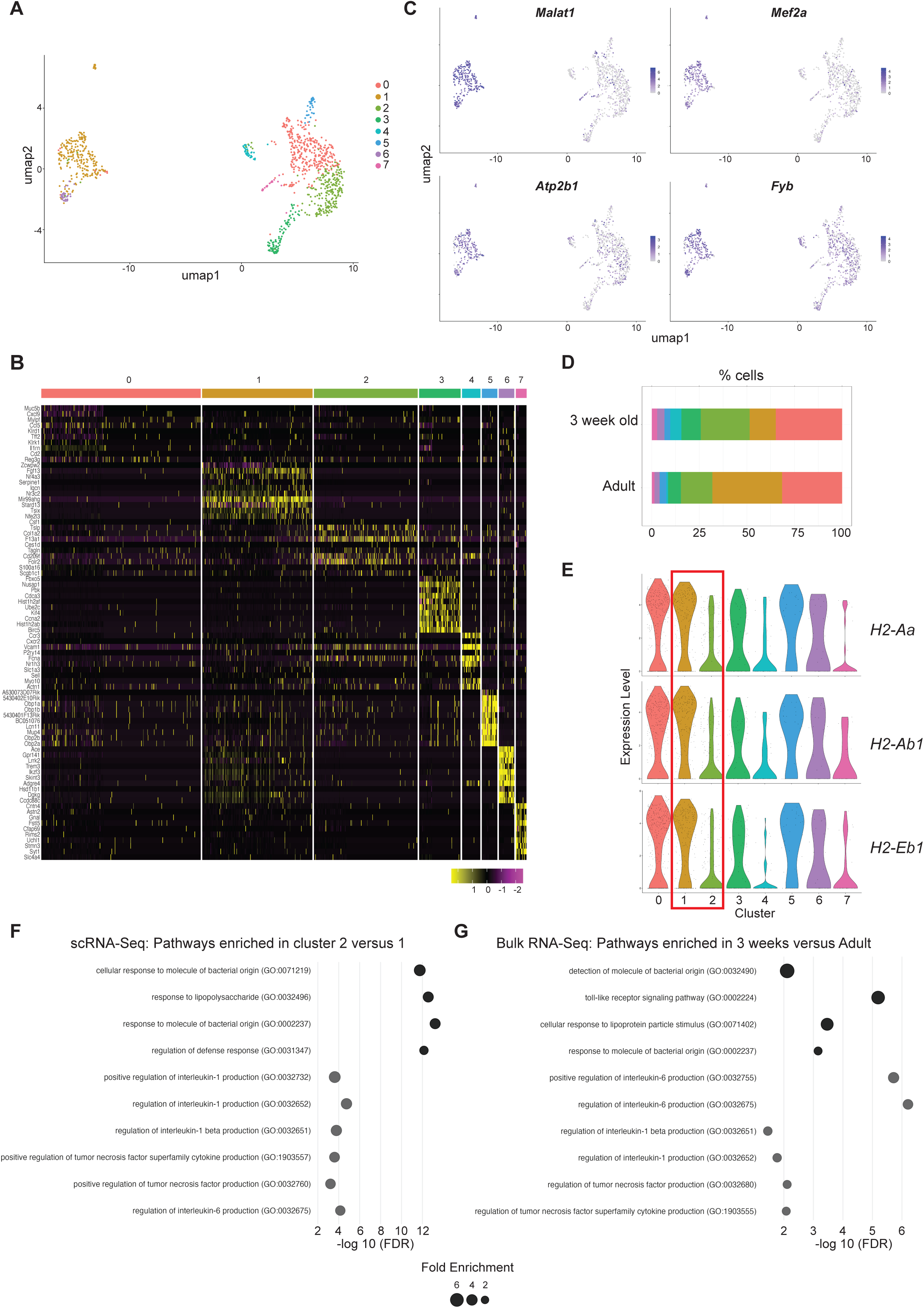
scRNA-sequencing of gingiva macrophages reveals that at 3 weeks of age there is an increase in macrophages enriched in bacterial response genes. Single-cell RNA-sequencing was undertaken on macrophages (CD45^+^Lin^-^CD11b^+^Ly6C^low^CD64^hi^) FACS sorted from gingiva of 3-week old and adult mice. **(A)** Combined UMAP dimensionality reduction analysis of cells from scRNA-seq of total sorted gingiva macrophages. **(B)** Heatmap showing the top 10 most DEGs by each cluster defined in (A). **(C)** UMAP showing expression of indicated genes across gingiva macrophage clusters. **(D)** Stacked bar graph showing proportions of cells in each gingiva macrophage cluster at 3 weeks and in adults. **(E)** Violin plots showing expression of *H2-Aa, H2-Ab1* and *H2-Eb1* genes in macrophage clusters. **(F)** DEGs were generated comparing clusters 1 and 2 and GO terms associated with these DEGs shown for pathways associated with Bacterial response (black) and Cytokine response (grey). **(G)** Bulk-RNA-sequencing was undertaken on total macrophages FACS sorted from the gingiva of 3-week old and adult mice. DEGs were determined and GO terms associated with these DEGs shown for pathways associated with Bacterial response (black) and Cytokine response (grey).

All macrophage clusters were present in 3 week old and adult gingiva (**Fig. S4D**). However, the relative representation of each cluster was slightly different at 3 weeks compared to adult gingiva (**Fig. 4D**), with more gingiva macrophages in cluster 2 at 3 weeks (light green) and more in cluster 1 in adults (mustard). In line with the expansion of MHCII^-^gingiva macrophages seen at 3 weeks by flow cytometry, cluster 1 was enriched for MHCII genes compared to cluster 2 (**Fig. 4E**). Comparing cluster 1 and 2 directly, there were a number of differentially expressed genes which could be used to examine enrichment of gene pathways. Cluster 2, which was increased in the gingiva of 3 week old mice, was highly enriched in genes associated with response to bacteria and bacterial-derived products (**Fig. 4F**). Also enriched, although to a lesser extent, were pathways associated with IL-1, IL-6 and TNF production (**Fig. 4F**). This enrichment of gene pathways indicating responses to bacteria and pro-inflammatory cytokine production in macrophages in the gingiva of 3 week old mice, could also be seen in bulk RNA-sequencing of total macrophages sorted from 3 week old compared to adult mice (**Fig. 4G**). These data suggest an expanded population of macrophages within the gingiva at 3 weeks of age that represents a population of activated macrophages responding to microbial stimuli.

### Early-life increases in gingiva macrophages are required to restrict oral commensal bacterial biomass

Based upon our RNA-sequencing data we next explored the functionality of macrophages in the gingiva of 3 week old and adult mice. Macrophage uptake of pHrodo beads and production of ROS were similar when comparing those from 3 week old and adult gingiva, and also comparing MHCII^-^ and MHCII^+^ cells (**Fig 5A,B**). As macrophage numbers were increased in 3 week old animals compared to adults, the total number of gingiva macrophages either phagocytosing or producing ROS were increased in 3 week old mice compared to adults, but only within the MHCII^-^population (**Fig. S5A**). However, upon *ex vivo* stimulation with LPS, proportions of macrophages staining positive for TNFα and IL-1β, but not IL-1α, were increased in 3 week old gingiva compared to adult, specifically within the MHCII^-^macrophage population (**Fig. 5C**). These data indicate that the increased MHCII^-^macrophage population present in the gingiva at 3 weeks of age exhibited an elevated capacity to produce inflammatory cytokines.

**Figure 5:**
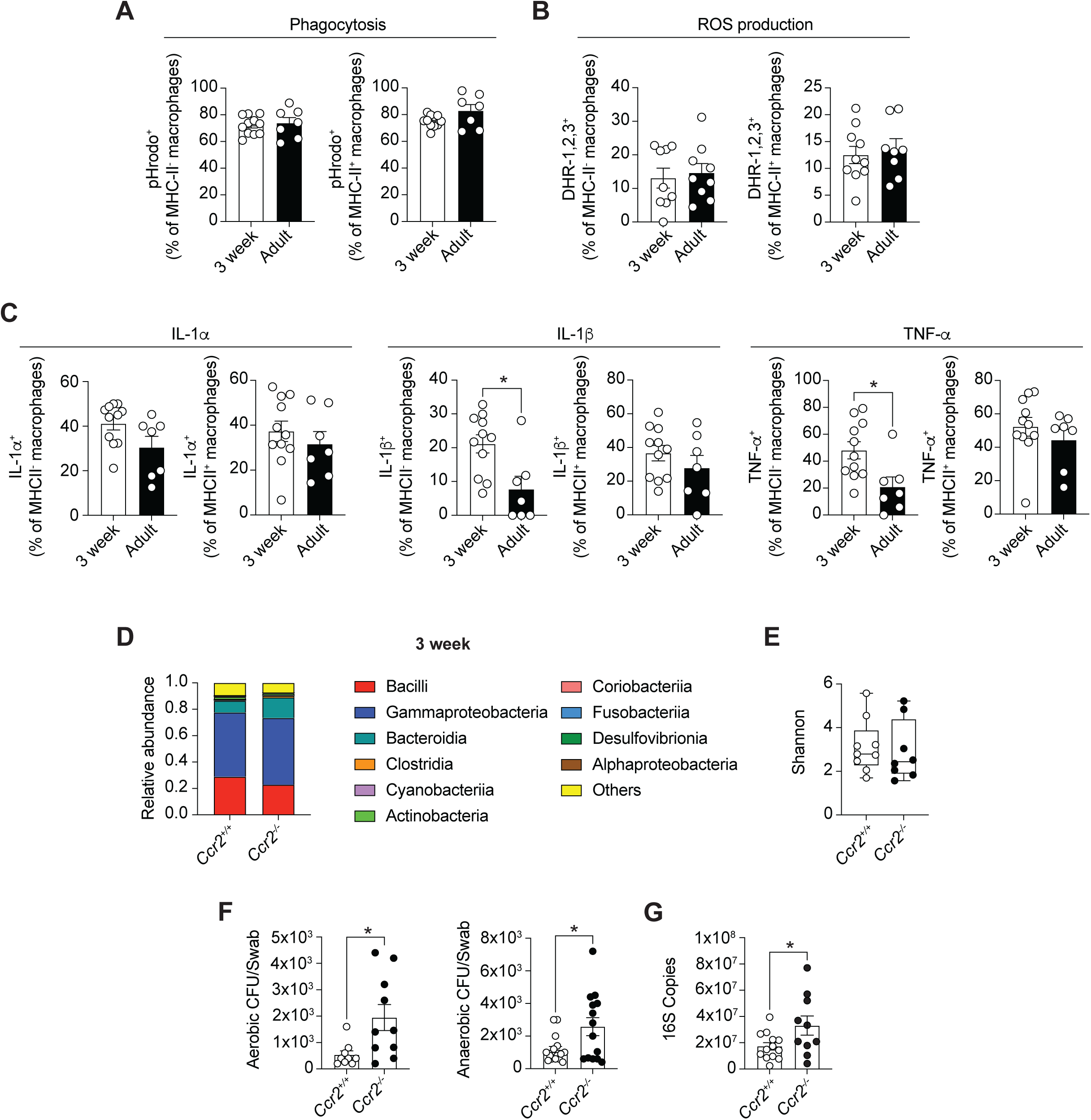
Increases in gingiva macrophages at 3 weeks of age are vital to restrict the biomass of the oral microbiome. **(A)** Frequency of pHrodo^+^ MHC-II^+^ and MHC-II^-^gingival macrophages in 3-wk and 8-10-wk-old C57BL/6 WT mice. **(B)** Frequency of DHR-1,2,3^+^ MHC-II^+^ and MHC-II^-^gingival macrophages in 3-wk and 8-10-wk-old C57BL/6 WT mice. **(C)** Frequency of IL-1α^+^ (left), IL-1β^+^ (centre) and TNF-α^+^ (right) MHC-II^+^ and MHC-II^-^gingival macrophages in 3-wk and 8-10-wk old C57BL/6 WT mice in response to LPS stimulation. **(D)** Relative abundance of bacterial species isolated from oral swabs of 3-wk-old *Ccr2^+/+^* and *Ccr2^−/−^*mice. **(E)** Shannon diversity index of bacterial species isolated from oral swabs of 3-wk-old *Ccr2^+/+^* and *Ccr2^−/−^* mice. **(F)** Aerobic (left) and anaerobic (right) colony forming units cultured from oral swabs of 3-wk-old *Ccr2^+/+^* and *Ccr2^−/−^*mice. **(G)** 16S copies as determined by qPCR of DNA isolated from oral swabs of 3-wk-old *Ccr2^+/+^* and *Ccr2^−/−^* mice. Data are representative of at least two individual experiments, n=3-5 per experiment. Error bars show mean±SEM. Statistical comparisons were performed by two-way ANOVA (A, B, C) or Welch’s *t* test (F,G): *P≤0.05.

Thus far we have shown that over an early life window of 3 weeks, numbers of gingiva macrophages increase in response to microbial signals. Examination of macrophages in the gingiva at 3 weeks of age demonstrated that these macrophages are enriched in genes associated with responding to bacteria, TNFα and IL-1β production. Our functional assays show macrophages from the gingiva of 3 week old mice were more responsive to LPS than adult gingiva macrophages. To understand the broader implications of this early life recruitment and activity of gingiva macrophages, we examined the oral commensal bacterial community, hypothesizing that the transient macrophage expansion at 3 weeks was vital to curate this diverse bacterial community. As we have demonstrated that transient expansion of macrophages at 3 weeks of age was dependent on CCR2 (**Fig. 3G,H**), we compared the oral microbiome of control and *Ccr2^−/−^* mice at 3 weeks of age. Neonates examined were littermates and remained cohoused until examined at 3 weeks of age. Failure to expand the gingival macrophage compartment at 3 weeks of age had no impact on the bacterial community structure, with both control and *Ccr2^−/−^* mice exhibiting very similar commensal bacterial communities (**Fig. 5D,E**). However, failure to expand gingiva monocyte-derived macrophages at 3 weeks of age did result in an elevated oral bacterial load, with the bacterial biomass, as determined by both CFU and 16S copy number, increased in 3 week old *Ccr2^−/−^* mice compared to littermate controls (**Fig. 5F,G**). This elevation in oral bacterial load was only observed in 3 week old *Ccr2^−/−^*animals, as by adulthood, when gingiva macrophages persist in lower frequencies, the oral bacteria biomass was similar between littermate controls and *Ccr2^−/−^* mice (**Fig. S5B**). However, in adult mice the oral bacterial communities evolved differently and by this time point community composition was distinct (**Fig. S5C-E**). Together, these findings characterise the macrophage populations residing within healthy gingiva and demonstrate that at early time-points after birth macrophages within this barrier play key roles in limiting microbial colonisation.

## Discussion

Substantive research effort has defined macrophage ontogeny and function in most organs, yet the development, turnover and activities of gingival macrophages remained minimally explored. Here we delineated the ontogeny of gingiva macrophages, outlining that although embryonic progenitors initially seed the gingival macrophage compartment, these are displaced by monocyte-derived macrophages. Interestingly, these monocyte-derived macrophages do not differentiate into long-lived, self-renewing cells. Conserved across other barrier tissues are macrophage populations that self-renew; one key difference in the gingiva was that we could not identify a population of long-lived macrophages. Despite a small subset of proliferative macrophages identified in the gingiva via our scRNA-sequencing data, fate-mapping approaches indicated few macrophages persisted long-term. A previously established paradigm, suggested that macrophages in the tonically inflamed GI tract could not persist for long periods of time [15]. Subsequently, we and others demonstrated that this was not the case and there is a population of resident macrophages (many of which are delineated by expression of Tim-4 [11, 16]). Our data here would indicate that the situation for macrophage turnover in the gingiva is more akin to that originally proposed for the gut; but what could drive this consistent replenishment of macrophages within the gingiva of adult mice? The gingiva is exposed to a diverse and dense bacterial commensal community, a stimulus that contributes to macrophage replenishment rates at other sites. However, this exposure did not drive the high level of replenishment seen in the gingiva, as reduction in oral bacterial load did not impact macrophage replenishment rates. Given the tissue damage continuously experienced at the gingiva concomitantly with microbial stimuli, it is possible these challenges combined could curtail gingiva macrophage lifespan and drive continuous replenishment. In line with this, continuous replenishment of Langerhans cells in the gingiva has been shown [18], contrasting the slower turnover of Langerhans cells in the skin, further highlighting the high levels of challenge experienced at the gingiva.

Alongside exhibiting elevated replenishment rates, we show that over the neonatal period from 1-4 weeks the gingiva macrophage network was substantially remodelled. We found that the total number of gingiva macrophages was increased at 3 weeks of age. This was preceded by an increase in monocytes just after 2 weeks post birth and, moreover, this increase in macrophages did not occur in 3 week old *Ccr2*-deficient animals. Thus, we detail a temporally restricted recruitment of monocytes and subsequent macrophage development in the gingiva. Previous work has shown a gradual increase in the total number of immune cells in the gingiva over the weaning period into adulthood [46]. In contrast, the increase in macrophages at 3 weeks of age was transient and by 4 weeks post birth the number of macrophages had fallen to a level at which it then persisted into adulthood. Alongside the number of macrophages changing over the neonatal period, macrophage phenotype changed too. Immediately post birth, most macrophages in the gingiva were Tim4^+^. Yet, as the macrophage population expands and then contracts over the neonatal period, this Tim4^+^ population was largely lost. Barrier immune network remodelling is common over the neonatal period, with increased recruitment of monocytes, and subsequent monocyte-to-macrophage differentiation, shown to occur to meet the demands of newly forming tissue structures [39, 40, 47]. Our data outline a time-restricted expansion of gingiva macrophages, suggesting not only a specific requirement for macrophages at this time-point, but also a developmentally restricted gingiva niche that is open and into which macrophages can differentiate specifically during this neonatal window. If such a niche exists, then the specific factors supporting differentiation into this niche remain to be determined, but we show that a key stimulus for entry into this niche is dependent on the microbiota. Our data clearly show that in 3 week old germ-free animals, numbers of gingiva macrophages are not increased. Thus, despite the oral microbiome not being the key driver for gingiva macrophage replenishment in adults, this local microbial community plays a key role in shaping the gingiva macrophage network in early life.

Microbial cues experienced in early life significantly impact barrier immune networks, recruiting and educating immune cells to safeguard effective host-commensal interactions [48]. In the gingiva, over the neonatal period, colonising microbiota have been shown to induce neutrophil, Langerhans cell and γδ T cell recruitment, plus antimicrobial peptide expression [46, 49, 50]. Here we tie microbe-induced expansion of macrophages in the gingiva, to local control of the microbial community, highlighting a key role for recruited macrophages in limiting the biomass of the oral commensal community. Exploration of the function of gingiva macrophages from 3 week old and adult animals *ex vivo* demonstrated macrophages from the gingiva of 3 week old mice exhibited elevated pro-inflammatory cytokine production in response to LPS stimulation, exhibiting enhanced production of both TNFα and IL-1β. Indeed, our RNA-sequencing approaches also showed enrichment of gene pathways involved in production of both these cytokines, alongside IL-6, in neonatal versus adult gingiva macrophages. Are neonatal macrophages simply more activated, as shown for other sites [51]? Perhaps, but adult gingiva macrophages still produce high levels of cytokines, unlike gut macrophages in which production of TNFα is attenuated [3]. This hyporesponsive programme of gut macrophages is driven by dwell time in the tissue and therefore unlikely to occur in the gingiva where we show macrophages are shorter-lived. Nevertheless, early-life macrophage production of TNFα, IL-1β, and/or IL-6 could well co-ordinate gingiva immune responses to constrain local microbes. Indeed, gut macrophages act as central immune hubs producing IL-1β to restrict local microbes [1]. In the gingiva, IL-1β could also be acting as a key mediator of oral microbiome curation during this neonatal window.

Here we also undertook scRNA-sequencing of gingiva macrophages in both adult and 3 week old mice to explore cellular heterogeneity. We defined two broad populations of macrophages in the gingiva, with one expressing higher levels of M1-associated proinflammatory genes (population A). The second, population B, was more diverse yet it was possible these two discreet populations represent macrophages in distinct niches and/or which have distinct origins. Considering niche, our imaging shows that in adult gingiva monocytes (IBA^+^F4/80^-^) localise to different regions; both the submucosa and epithelium. In contrast, macrophages (IBA^+^F4/80^+^) are only seen within the submucosa. Our recent work has shown that haematopoietic stem and progenitor cells (HSPC) are resident within the gingiva in health [30]; it is possible that the two macrophage populations represent those derived from local versus distal haematopoiesis. Beyond this, gingiva macrophages were defined as 8 discreet clusters. Of the remaining clusters it was difficult to assign specific signatures associated with other RTM populations but, given the macrophage populations defined within other barriers, it will be intriguing to further explore whether the gingiva contains a population of neurone-associated macrophages, especially considering the glial cell network resident within this tissue [52]. Moreover, given the high levels of damage experienced at the gingiva, it will also be important to better understand whether and how discreet macrophages populations contribute to healing at this site. Indeed, a previous study outlined a population of CD74^+^ macrophages which emerged during resolution of periodontitis [23], however this was not a subset we identified in our dataset. Intriguingly, despite the elevated recruitment of monocytes to the gingiva at 2 weeks, and subsequent expansion of gingiva macrophages at 3 weeks, all macrophage clusters were identified in both adult and 3 week old gingiva. Indeed, the only evident change was the balance between cluster 1 and 2, with both single-cell and bulk RNA-sequencing datasets supporting an enrichment of bacterial responses gene signatures and cytokine production pathways in macrophages from 3 week old mice compared to adults. This was in line with our functional data which showed macrophages from gingiva of 3 week old mice exhibited elevated responses to LPS compared to macrophages from adult gingiva.

Combined, our results demonstrate that gingiva macrophages are continuously replenished as, unlike within other barriers, we were unable to identify a long-lived population in the gingiva. This constant replenishment is not driven by the high microbial burden at this site, but the elevated turnover rates could well place gingiva macrophages at the top of a barrier macrophage spectrum, above the gut, denoting them globally as the tissue macrophage population with the lowest self-renewing capacity. Although microbe-induced signalling did not condition macrophage replenishment, these signals did play key roles in re-shaping the macrophage network in early-life. We show that microbial-signals drive early-life increases in gingiva macrophages which are transient but vital to restrict the growth of the oral commensal community. How important this transient increase in gingiva macrophages is for long-term tissue health remains to be explored, but similar to other weaning reactions described across barrier sites, it is possible that failure to establish effective host-microbe dialog in early life could have later life inflammatory consequences. Our findings provide a new perspective on macrophages within the gingiva and, as gene pathways enriched in macrophages at 3 weeks of age are also ones that become dysregulated in the context of periodontitis, our work provides a foundation from which the therapeutic potentials of these cells can be explored.

## Materials and methods

### Mice

Mice aged between 1-12 weeks were housed in individually ventilated cages, under SPF conditions, with standard chow and water provided *ad libitium*, at a constant temperature and a 12h light/dark cycle. C57BL/6J mice were purchased from Envigo. *Cx3cr1^CreER^xR26-yfp* (bred from *Cx3cr1^CreER^* and *R26R-yfp* strains from The Jackson Laboratory; [10, 53]*), Ccr2^−/−^*(originally from The Jackson Laboratory;[54]), *Ms4a3^TdT^ (*bred from B6.Ms4a3em2(cre)Fgnx [12] and Gt(ROSA)26Sortm14(CAG-tdTomato)Hze strains from the Shanghai Model Organisms Centre and The Jackson Laboratory respectively) and *Cdh5*-CreERT2:Rosa^tdT^ (kind gift from Rebecca Gentek [38]) were bred in-house. Mice were age- and sex-matched within experiments. All experiments were approved by local Animal Welfare Ethical Review Body and the Home Office, UK.

### Tamoxifen treatment

For administration to adult *Cx3cr1^CreER^xR26-yfp* mice, tamoxifen (Sigma-Aldrich) was dissolved in 10% ethanol and 90% corn oil to a concentration of 100 mg/ml. Mice were dosed with 5mg by oral gavage for five consecutive days. For *Cdh5*-CreERT2 fate mapping, dosing was performed as previously described [38].

### Antibiotics

*Cx3cr1^CreER^xR26-yfp* mice received an antibiotic cocktail containing ampicillin, metronidazole, neomycin, gentamicin (all 1g/L) and vancomycin (0.5g/L) in the drinking water starting the day after the final dose of tamoxifen and continuing for 4 weeks.

### Preparation of single-cell suspensions

#### Gingiva

Gingiva were dissected as previously described [55]. Tissue blocks were digested for 30 min a 37°C in serum-free medium containing 3.2 mg/ml collagenase IV (Gibco) and 7.5 μg/ml DNase. Following removal of gingiva tissue from maxilla and mandible, single-cell suspensions were obtained by mashing tissues through a 70 μm strainer.

#### Small intestine

Cells were isolated as previously described [11].

#### Bone marrow

Femurs were flushed, and red blood cell lysis performed using ACK lysing buffer to obtain single cell suspensions.

#### Brains

Brains were homogenised using a Dounce homogeniser. Homogenate was passed through a 70 μm strainer and centrifuged. The supernatant was discarded, the pellet resuspended and centrifuged on a Percoll gradient. The uppermost myelin layer was removed and remaining pellet washed in PBS and resuspended to obtain a single cell suspension.

#### Skin

Dorsal and ventral halves of ears were separated. Samples were digested for 1hr 45 min at 37°C with agitation in serum-free medium containing 0.25 mg/ml LIberase TL (Roche) and 0.5 mg/ml DNase. Tissue was further processed using a MediMachine (BD Biosciences) for 6 min. Suspensions were passed over a 100 μm strainer to obtain single cell suspensions.

### Flow cytometry

Single-cell suspensions from tissues were washed in PBS and incubated with antibody cocktails plus anti-CD16/32 (2.4G2; BioXcell) for 15-20 min at 4°C. Cells were stained with combinations of the following antibodies, which were obtained from eBioscience and BioLegend: CD45 (clone 30-F11), CD11b (clone M1/70), Ly6G (clone 1A8), CD3e (clone 145-2C11), B220 (clone RA3-6B2), NK1.1 (clone PK136), CD11c (clone N418), Ly6C (clone HK1.4), CD64 (clone X54-5/7.1), MerTK (clone 2B10C42), Tim-4 (clone RMT4-54), CX3CR1 (clone SA011F11), F4/80 (clone BM8), IBA1 (clone EPR16588), MHC-II (clone M5/114.15.2), CD163 (clone S15049l), Ki-67 (clone SolA15), IL-1α (clone ALF-161), IL-1β (clone NJTEN3), TNF-a (clone MP6-XT22), EpCAM (clone G8.8), CD207 (clone eBioL31). Dead cells were excluded by use of a Live/Dead Fixable blue dead cell stain kit (Molecular Probes). Following surface staining, cells were fixed in eBioscience Foxp3/Transcription Factor Fixation buffer for 20 min at 4°C. Samples were then stained overnight at 4°C with antibodies diluted in eBioscience Foxp3 permabilisation buffer. Samples were acquired using a BD LSRFortessa (BD Biosciences) flow cytometer and analysed with FlowJo (Treestar).

### Monocyte and macrophage isolation by FACS

Cells were sorted using a BD FACSAria Fusion (BD Biosciences). Monocytes and macrophages were sorted using the gating as in **Fig.1A**.

### Cytospin

Gingiva monocytes and macrophages FACS purified were mounted on Superfrost slides (Thermo Fisher Scientific) using a Cytospin centrifuge (Cytospin 4; Thermo Fisher Scientific). Cells were fixed with ice-cold methanol before staining with haematoxylin and acidic eosin (Thermo Fisher Scientific) and mounted with DPX (Thermo Fisher Scientific). Images were captured using an Olympus BX63 upright microscope, processed and analysed using FIJI2.

### Immunofluorescent imaging

Gingiva were fixed for 25h in 10% neutral buffered formalin before decalcification in 14% EDTA for 14 days. Tissues were then incubated in 30% sucrose overnight before embedding in OCT and frozen on dry ice. Sections were incubated in blocking buffer containing 0.1M Tris-HCl (Invitrogen, 10434742), 1:200 Triton-X 100 (Sigma-Aldrich, X100-5ML), 1:100 mouse serum (Sigma-Aldrich, M5905-5ML) and 1:100 rat serum (Sigma-Aldrich, R9759-5ML) and 1% BSA (R&D Systems, DY995) at room temperature for 1hr. Sections were then incubated with primary antibodies, 1:200 rabbit anti-Iba1 (Abcam, ab178846) and 1:200 anti-F4/80-Alexa Fluor 647 (Abcam, ab204467) in blocking buffer overnight at 4°C. Sections were washed three times in PBS for 10 minutes before incubation with secondary antibodies (donkey anti-rabbit Alexa Fluor 488, 1:250) at room temperature for 1hr. Sections were washed three times in PBS for 10 minutes before mounting in Fluoromount-G Mounting Medium (Invitrogen, 00-4958-02). Z-stack tile scans of gingiva sections were acquired on a Leica SP8 inverted confocal microscope (Wetzlar, Germany). Tile scans were acquired with 40x oil-immersion objective, 1X Zoom, with individual channels on separate track series, with an optimised Z-step size of 0.2-0.5µm. Images were analysed with Imaris software (Bitplane) and FIJI2.

### Single cell RNA-sequencing

#### Library Generation

Gene expression libraries were prepared from single cells using the Chromium Controller and Single Cell 3ʹ Reagent Kit v3.1 (10x Genomics, Inc. Pleasanton, USA) according to the manufacturer’s protocol (CG000315 revision E). Briefly, nanoliter-scale Gel Beads-in-emulsion (GEMs) were generated by combining barcoded Gel Beads, a master mix containing cells, and partitioning oil onto a Chromium chip. Cells were delivered at a limiting dilution, such that the majority (90-99%) of generated GEMs contained no cell, while the remainder largely contained a single cell. The Gel Beads were then dissolved, primers released, and any co-partitioned cells lysed. Primers containing an Illumina TruSeq Read 1 sequencing primer, a 16-nucleotide 10x Barcode, a 10-nucleotide unique molecular identifier (UMI) and a 30-nucleotide poly(dT) sequence were mixed with the cell lysate and a master mix containing reverse transcription (RT) reagents. Incubation of the GEMs yielded barcoded cDNA from poly-adenylated mRNA. Following incubation, GEMs were broken and pooled fractions recovered. First-strand cDNAwas then purified from the post GEM-RT reaction mixture using silane magnetic beads and amplified via PCR to generate sufficient mass for library construction. Enzymatic fragmentation and size selection were used to optimize the cDNA amplicon size. Illumina P5 and P7 sequences, i7 and i5 sample indexes, and TruSeq Read 2 sequence were added via end repair, A-tailing, adaptor ligation, and PCR to yield final Illumina-compatible sequencing libraries.

#### Sequencing

The resulting sequencing libraries comprised standard Illumina paired-end constructs flanked with P5 and P7 sequences. The 16 bp 10x Barcode and 12 bp UMI were encoded in Read 1, while Read 2 was used to sequence the cDNA fragment. Sample index sequences were incorporated as the i7 and i5 index reads. Paired-end sequencing (28:90) was performed on the Illumina NovaSeq6000 platform on 1343 cells at a read depth of ∼370764 reads per cell. The BCL sequence data were processed for QC purposes using bcl2fastq software (v. 2.20.0.422) and the resulting FASTQ files assessed using FastQC (v. 0.11.3), FastqScreen (v. 0.14.0) and FastqStrand (v. 1.13.1) prior to pre-processing with the CellRanger pipeline.

#### Data pre-processing

The sequence files generated from the instrument were then processed using 10x Genomics custom pipeline Cell Ranger v 7.1.0, which generated the FASTQ files, aligned those files to the required genome, identified the valid barcodes as cells, counted UMIs and reported a gene by cell count matrix in sparse matrix format. Sequences were mapped to the mm10 genome provided with the Cell Ranger package. Resultant single cell MEX files were integrated using Seurat’s CCA and then processed following the standard Seurat vignette. Pseudotime analysis was performed using Monocle3 (v. 1.4.25). Cluster specific differentially expressed genes were identified using MAST (v. 3.19). Consequent gene ontology lists and scores were generated via PANTHER GO (https://pantherdb.org/).

#### Data availability

The gingiva macrophage dataset is available via the accession: E-MTAB-17147.

#### Code availability

The code used to analyse single cell RNAseq data is available on GitHub (https://github.com/rhdaw/gingiva_monocyte_analysis/).

### Bulk RNA-sequencing

Total RNA from FACS-sorted macrophages was isolated using the Single Cell RNA Purification Kit (Norgen Biotek Corp.). RNA sequencing was done by Novogene; RNA was amplified and libraries generated using SMART Seq v4 Ultra Low Input RNA Kit and Novogene NGS RNA Library Prep Set (PT042). Libraries were sequenced with NovaSeq Xplus 25B lane with 15Gb PE150 data per sample.

### Bacterial 16S qPCR and sequencing

The murine oral cavity was sampled for 30s using sterile ultrafine swabs as previously described [42]. Bacterial DNA was then isolated from these swabs using the DNeasy Powersoil Kit (Qiagen). For 16S qPCR, DNA samples were amplified using the following primers for total 16S forward; 5′-ACTCCTACGGGAGGCAGCAGT-3′ and reverse; 5′-ATTACC GCGGCTGCTGGC-3′. 16S copy number was determined by running a standard curve with a known 16S copy number. For 16S sequencing, amplification generation, sequencing and bioinformatics analysis were undertaken by Novogene. Briefly, the V4 16S ribosomal RNA (rRNA) region was amplified and amplicons sequenced on an Illumina platform and 250-base-pair paired-end reads were generated. Data filtration and chimera removal were done using Fastq and Vsearch, and then amplicon sequence variants (ASVs) were generated from the raw data using the dada2 pipeline. Subsequently, by applying QIIME2’s classify-sklearn algorithm, a pretrained Naive Bayes classifier was used for species annotation of each ASV, utilizing the Silva 138.1 database. This allowed abundance tables at the level of kingdom, phyla, class, order, family, genus, and species to be obtained.

### Bacterial CFU assessment

The murine oral cavity was sampled as above with sterile swabs that were placed in 200 μl of sterile PBS. Swabs were vortexed prior to bacterial isolation and serial dilutions of resultant bacterial suspensions plated onto LB or Wilkins-Chalgren (both Sigma-Aldrich) agar plates.

### In vitro assays

#### pHrodo

Phagocytic capacity was assayed using the pHrodo BioParticles Phagocytosis Kit for Flow Cytometry (Invitrogen) as per the manufacturer’s instructions using single cell suspensions generated from gingiva tissue.

#### ROS

Single cell suspensions of gingiva tissue were incubated with 50 ng/ml DHR-1,2,3 (Sigma-Aldrich) for 15 min at 37C before addition of PMA at a final concentration of 25 ng/ml for a further 40 min.

#### LPS stimulation

Single cell suspensions of gingiva tissue were stimulated with 10 ng/ml LPS in the presence of brefeldin A for 3 h.

### Statistics

Statistical analyses were performed using Prism (GraphPad Software). Data are presented as mean±SEM. For comparisons between two groups, a Welch’s *t* test was applied, where more than two groups were compared, a one-way ANOVA or two-way ANOVA was used as appropriate. Statistical significance was defined as P≤0.05.

## Acknowledgements

We acknowledge the University of Manchester Biological Service, Flow Cytometry and Genomics Technologies Core Facilities for providing access to equipment and expertise instrumental in the completion of this research.

## Funding

This work was funded by a Wellcome Trust Fellowship (224467/Z/21/Z) and a Lister Institute for Preventative Medicine Prize fellowship, both to JEK. Additional support was from a Kennedy Trust for Rheumatology Research Senior Fellowship to J.R.G and BBSRC grant BB/S01103X/2 to TNS. This work was also supported by the Medical Research Council Centre of Research Excellence in Exposome Immunology (UKRI/MR/B000935/1). Open Access funding provided by University of Manchester.

## Data/material availability

Requests should be directed to JEK.

## Author Contributions

Conceptualisation: JEK and KW; Methodology: KW, RD, DP, JC, FAM, SK, IEP, ZL, FG, RG, TNS, JRG, JEK; Investigation and data analysis: KW, RD, DP, JC, FAM, SK, AW, HMB, TC, IEP, RG, TNS, JEK; Writing-original draft: JEK, KW and JRG; Writing-review and editing: JEK, KW, TNS, FG, ZL and JRG; Funding acquisition: JEK and JRG; Supervision: JEK, KW, JRG, TNS and RG.

## Competing Interest Statement

The authors declare that there are no competing interests.

## Classification

Biological Sciences, Immunology and Inflammation

## This PDF file includes

Main Text Figures 1 to 5

## Supplemental Figures

**Figure S1:**
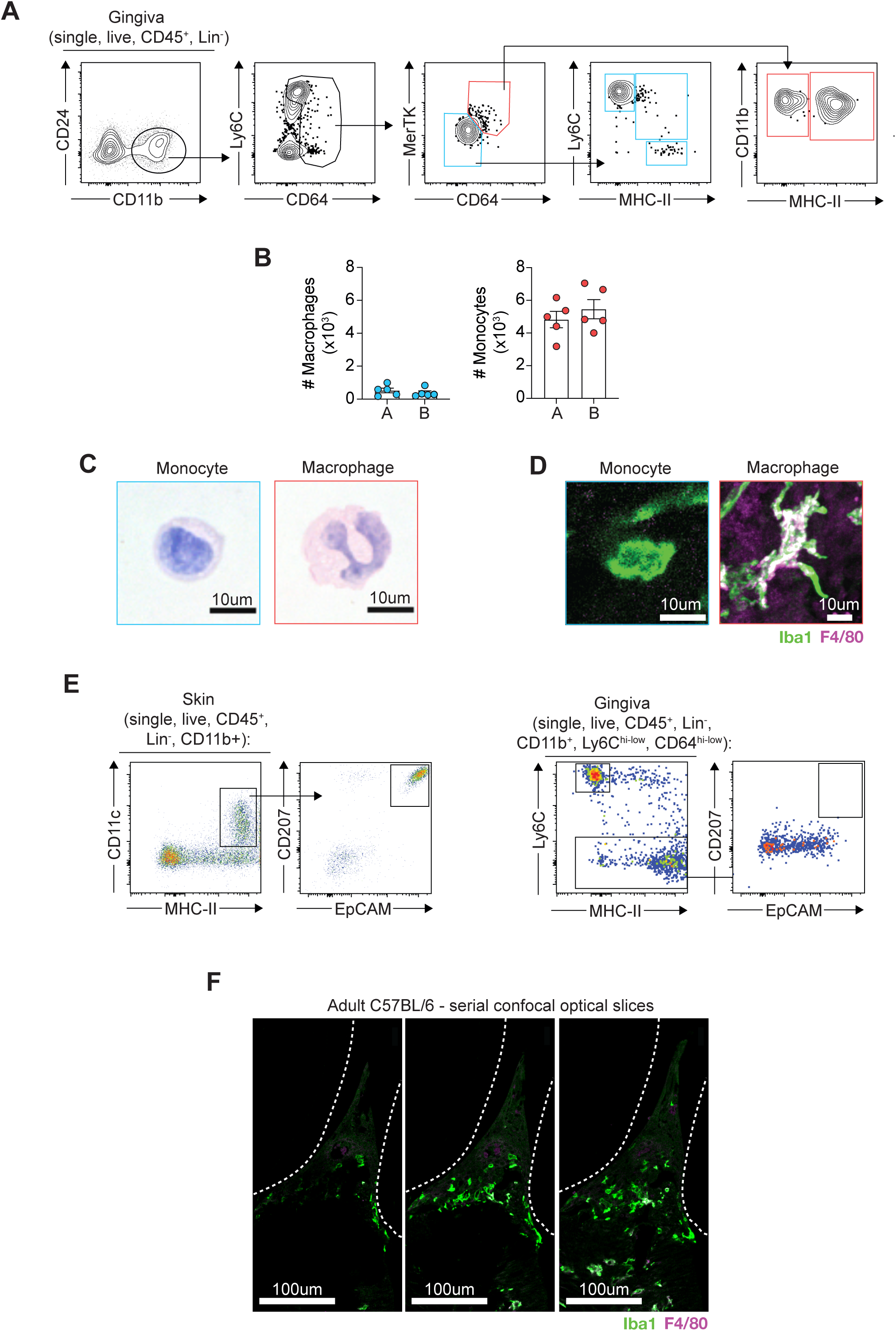
Gingiva macrophages are continuously replenished. **(A)** Alternative gating strategy used to identify monocytes and macrophage populations isolated from the gingiva. **(B)** Absolute numbers of gingival monocytes and macrophages as identified via gating strategy A (Figure 1A) and B (Figure S1 A). **(C)** Morphological characteristics as assessed by H&E staining of Ly6C^hi^ monocytes and macrophages sorted by FACS from the gingiva of 8-10-wk-old C57BL/6 WT mice. **(D)** Con-focal immunofluorescence microscopy of Iba1 and F4/80 in gingiva of C57BL/6 WT mice. **(E)** Representative flow cytometry plots showing identification of Langerhans Cells in the skin (left) and gingiva (right) based on expression of CD11c, MHC-II, CD207 and EpCAM. **(F)** Individual confocal optical sections of Iba1 and F4/80 in gingiva of adult C57BL/6 WT mice. White dotted line indicates outline of tooth. Data are representative of at least two individual experiments, n=2-3 per experiment. Error bars show mean ± SEM.

**Figure S2:**
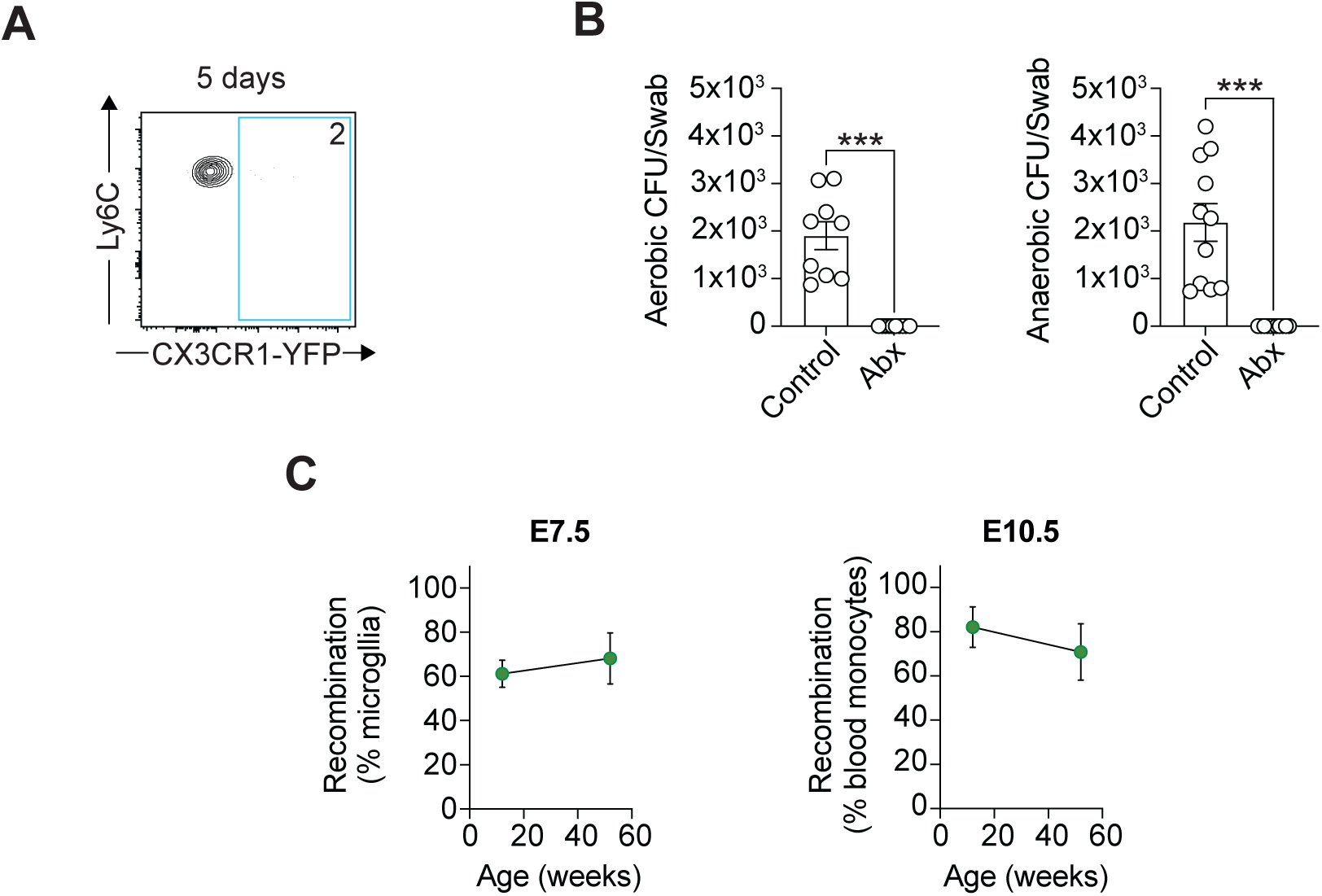
Transgenic strategies to track gingiva macrophage ontogeny and turnover. **(A)** Representative flow cytometry plot showing the frequency of YFP-expressing gingiva monocytes in *Cx3cr1^CreER^xR26-yfp* mice 5 d after tamoxifen treatment. **(B)** Aerobic (left) and anaerobic (right) colony forming units cultured from oral swabs of control and antibiotic (abx) treated animals. **(C)** Frequencies of reporter recombination in microglia (left) and blood monocytes (right) of *Cdh5*-CreERT2:Rosa^tdT^ mice pulsed with 4OHT at E7.5 and E10.5 respectively. Data are representative of at least two individual experiments, n=3-5 per experiment. Error bars show mean ± SEM.

**Figure S3:**
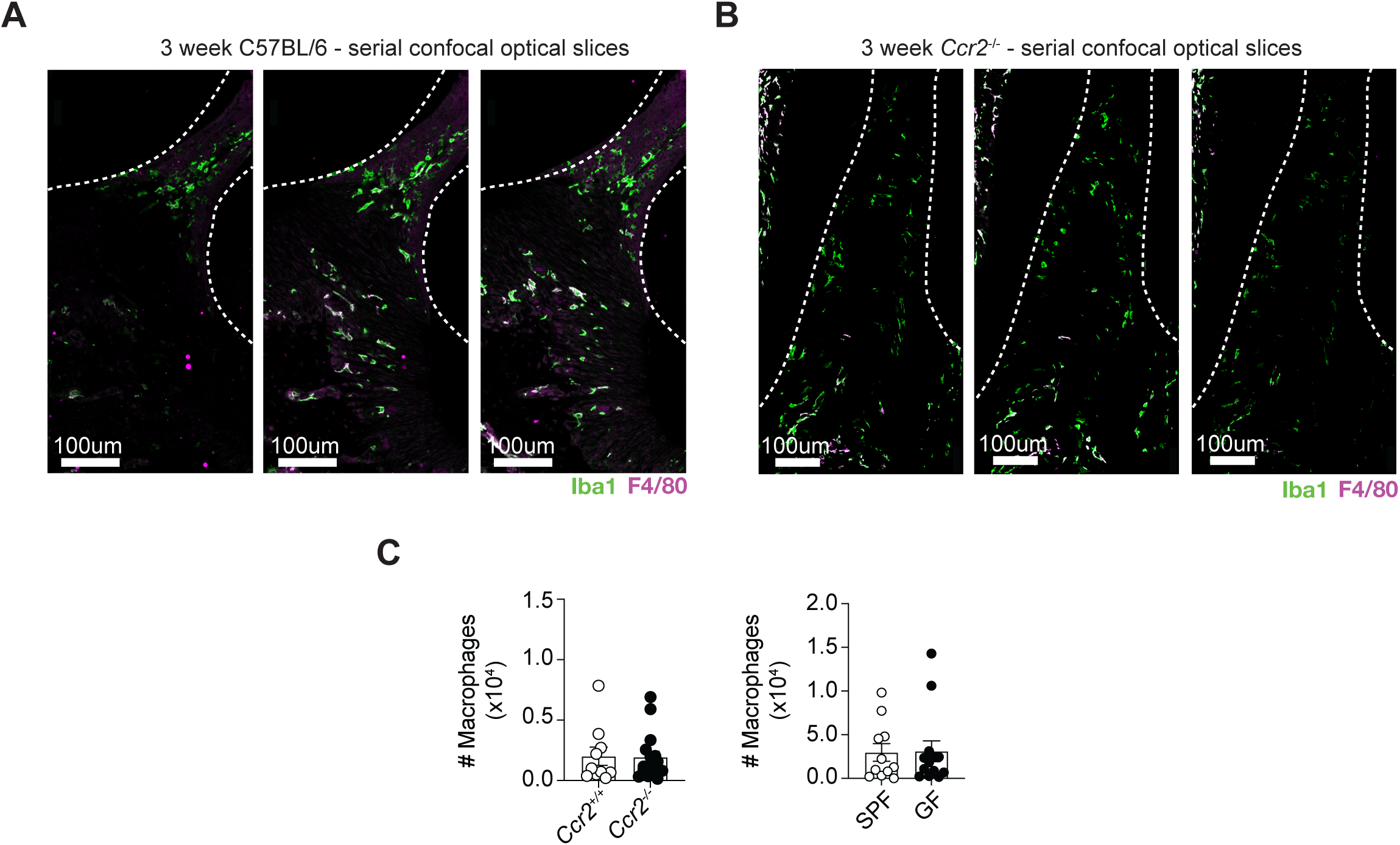
Adult *Ccr2*^−/−^ and Germ Free mice exhibit unaltered macrophage numbers. **(A)** Individual confocal optical sections of Iba1 and F4/80 in gingiva of 3-week old C57BL/6 WT mice. White dotted line indicates outline of tooth. **(B)** Individual confocal optical sections of Iba1 and F4/80 in gingiva of 3-week old *Ccr2*^−/−^ mice. White dotted line indicates outline of tooth. **(C)** Graphs show number of gingiva macrophages in adult *Ccr2*^+/+^ and *Ccr2*^−/−^ mice (left) and SPF and GF mice (right). Data are representative of at least two individual experiments, n=3-5 per experiment. Error bars show mean ± SEM.

**Figure S4:**
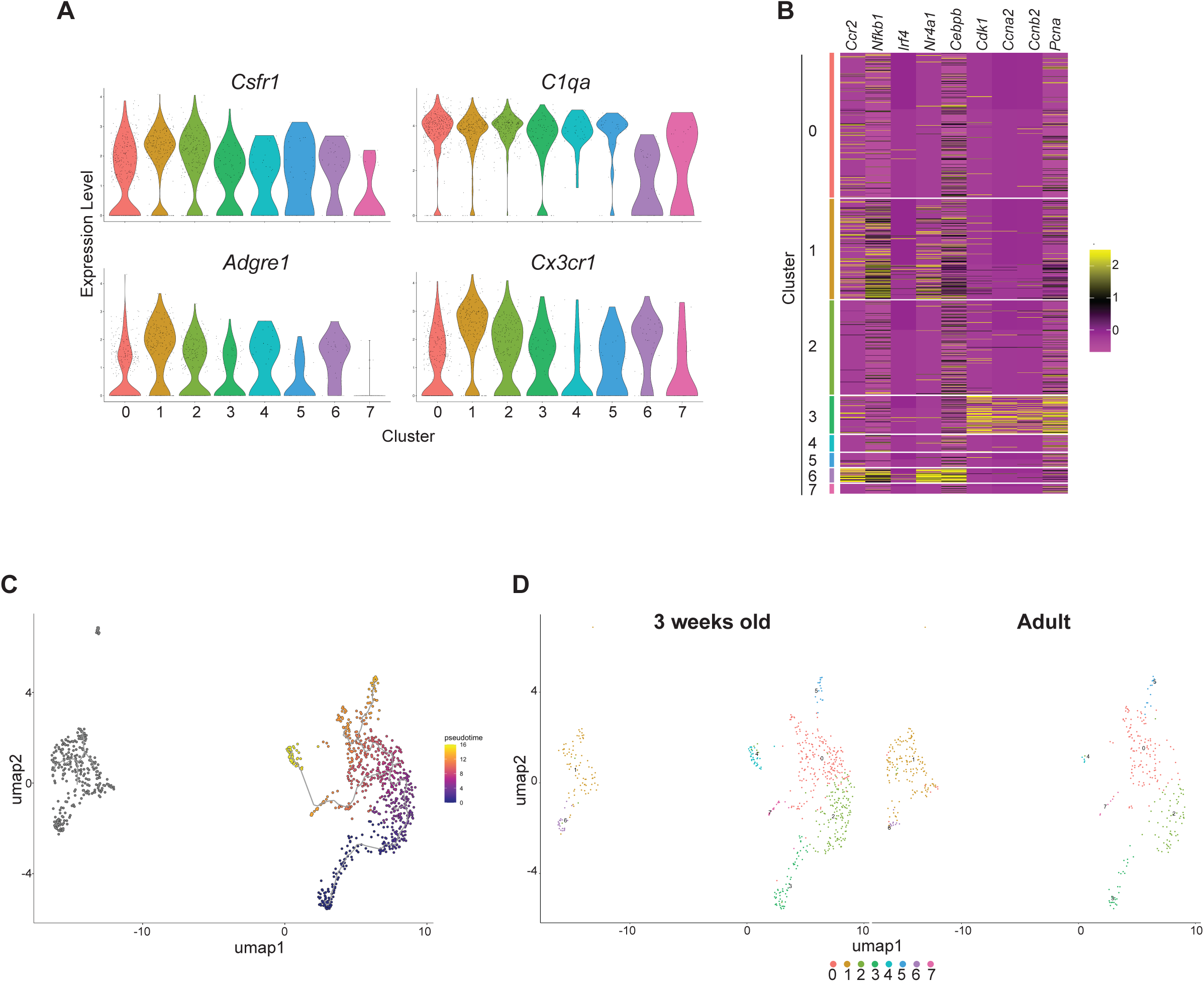
scRNA-sequencing of gingiva macrophages reveals 8 distinct clusters. Gingiva macrophages from 3 week old and adult animals were FACS-sorted and underwent scRNA-sequencing. **(A)** Violin plots showing gene expression of macrophage-defining genes. **(B)** Heat map showing expression of indicated genes in each macrophage cluster. **(C)** RNA velocity pseudotime was projected onto the UMAP shown in Fig. 4A. Dark blue marks the starting point and yellow the end points of the trajectory. **(D)** UMAP of cells from scRNA-seq of sorted macrophages from gingiva of 3 week old (Left) and adult (Right) mice.

**Figure S5:**
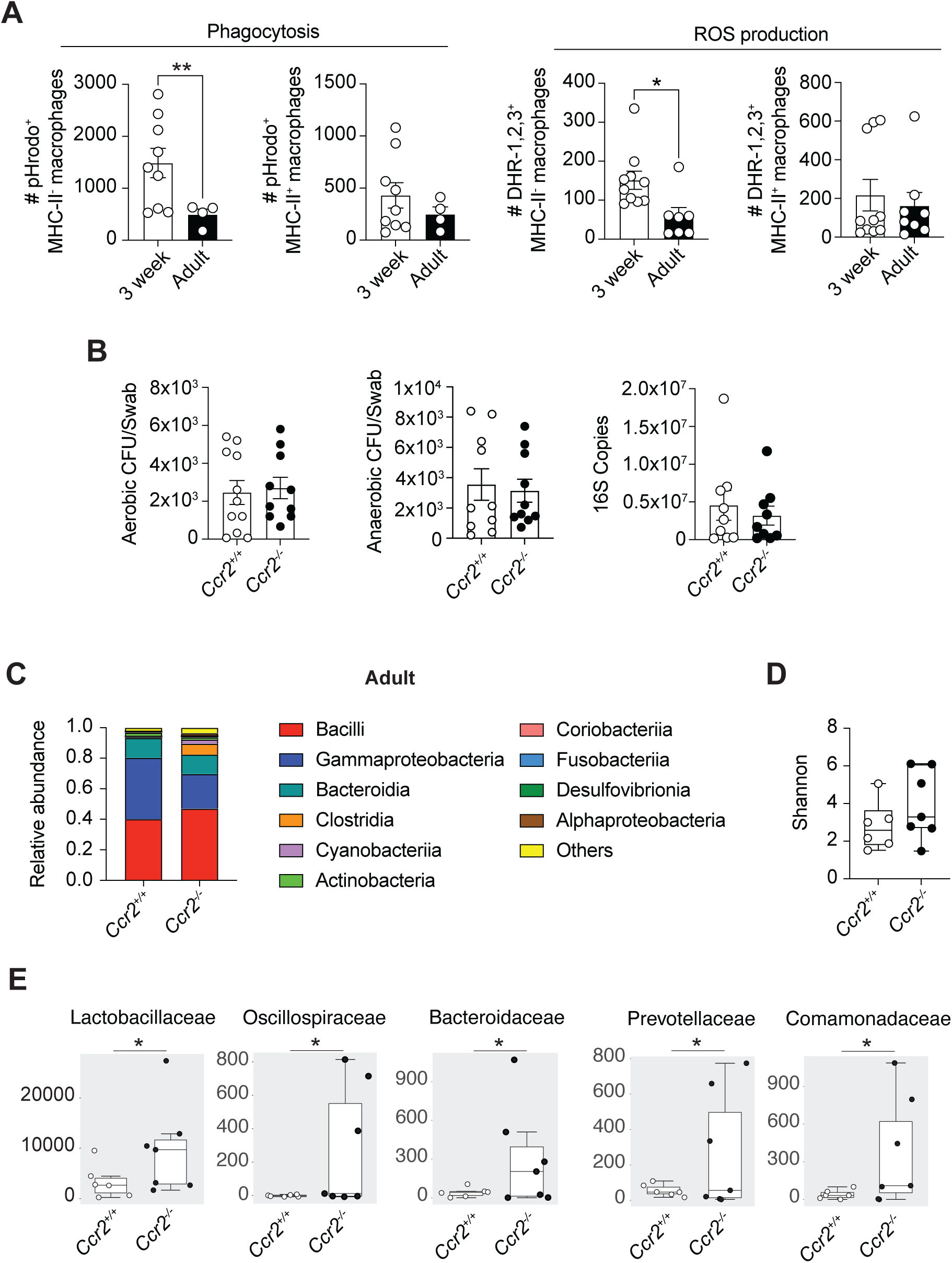
Functional activities of gingiva macrophages in 3 week old animals. **(A)** Total number of DHR-1,2,3^+^ and pHrodo^+^ MHC-II^+^ and MHC-II^-^ gingival macrophages in 3-wk and 8-10-wk-old C57BL/6 WT mice. **(B)** Aerobic (left) and anaerobic (centre) colony forming units cultured from oral swabs of adult *Ccr2*^+/+^ and *Ccr2*^−/−^ mice. 16S copies as determined by qPCR of DNA isolated from from oral swabs of adult *Ccr2*^+/+^ and *Ccr2*^−/−^ mice (right). **(C)** Relative abundance of bacterial species isolated from oral swabs of adult *Ccr2*^+/+^ and *Ccr2*^−/−^ mice. **(D)** Shannon diversity index of bacterial species isolated from oral swabs of adult *Ccr2*^+/+^ and *Ccr2*^−/−^ mice. **(E)** Relative abundance of bacterial families in adult *Ccr2*^+/+^ and *Ccr2*^−/−^ mice. Data are representative of at least two individual experiments, n=3-5 per experiment. Error bars show mean ± SEM. Statistical comparisons were performed by two-way ANOVA (A): *, P ≤ 0.05, **, P ≤ 0.01.

## Notes

### Competing Interest Statement

The authors have declared no competing interest.

