## Supplementary Figures for "Distinct monocyte replenishment kinetics of macrophages at the oral barrier restrict early-life bacterial colonisation"

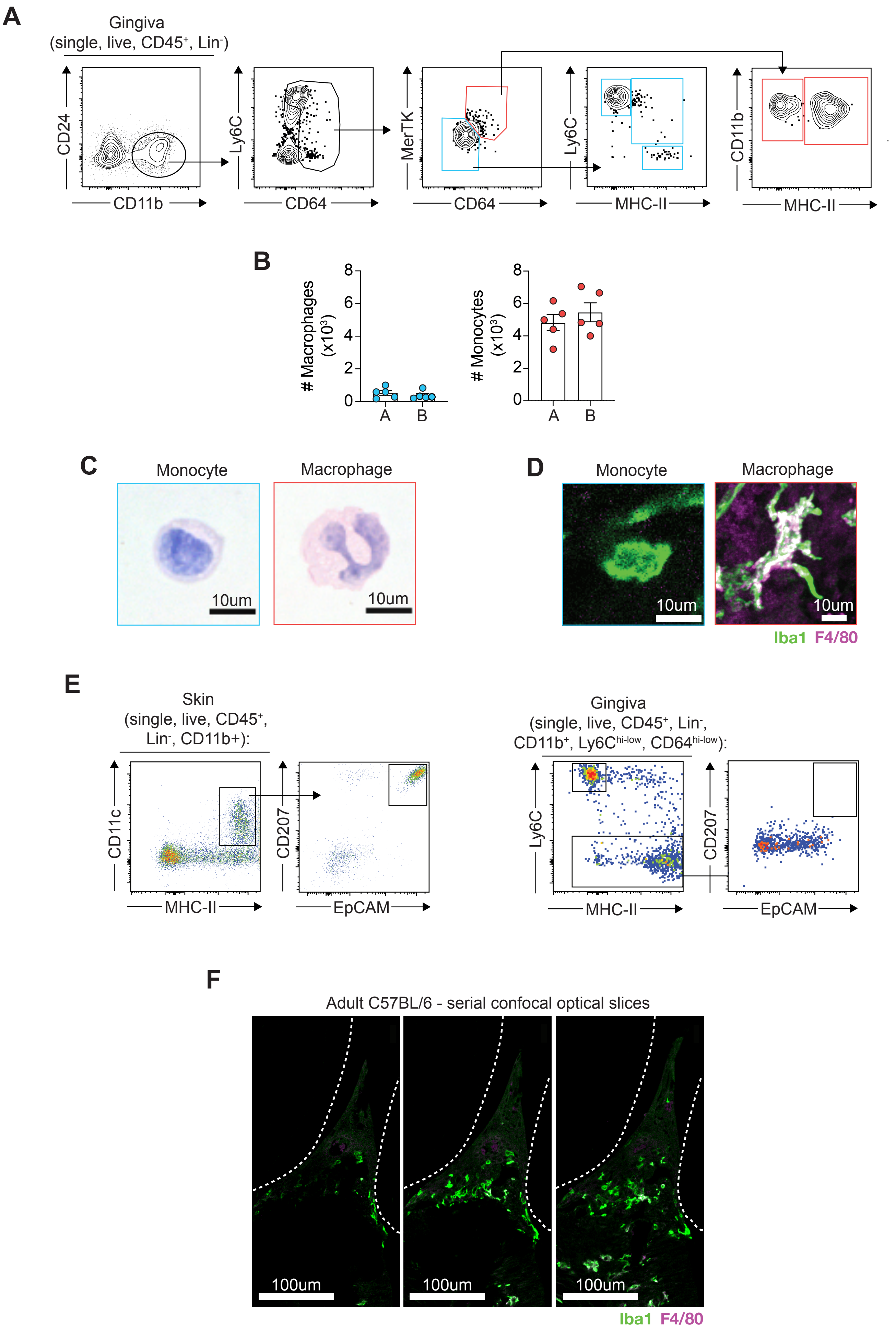

A
Gingiva
(single, live, CD45+, Lin-)
CD24
CD11b
Ly6C
CD64
MerTK
CD64
Ly6C
MHC-II
CD11b
MHC-II
B
### Macrophages (x10³)
### Monocytes (x10³)
0
2
4
6
8
A
B
C
Monocyte
Macrophage
10um
D
Monocyte
Macrophage
10um
Iba1 F4/80
E
Skin
(single, live, CD45+, Lin-, CD11b+):
CD11c
MHC-II
CD207
EpCAM
Gingiva
(single, live, CD45+, Lin-, CD11b+, Ly6Chi-low, CD64hi-low):
Ly6C
MHC-II
CD207
EpCAM
F
Adult C57BL/6 - serial confocal optical slices
100um
Iba1 F4/80

**Figure S1: Gingiva macrophages are continuously replenished**

**(A)** Alternative gating strategy used to identify monocytes and macrophage populations isolated from the gingiva. **(B)** Absolute numbers of gingival monocytes and macrophages as identified via gating strategy A (Figure 1A) and B (Figure S1 A). **(C)** Morphological characteristics as assessed by H&E staining of Ly6C$^{hi}$ monocytes and macrophages sorted by FACS from the gingiva of 8-10-wk-old C57BL/6 WT mice. **(D)** Confocal immunofluorescence microscopy of Iba1 and F4/80 in gingiva of C57BL/6 WT mice. **(E)** Representative flow cytometry plots showing identification of Langerhans Cells in the skin (left) and gingiva (right) based on expression of CD11c, MHC-II, CD207 and EpCAM. **(F)** Individual confocal optical sections of Iba1 and F4/80 in gingiva of adult C57BL/6 WT mice. White dotted line indicates outline of tooth. Data are representative of at least two individual experiments, n=2-3 per experiment. Error bars show mean ± SEM.

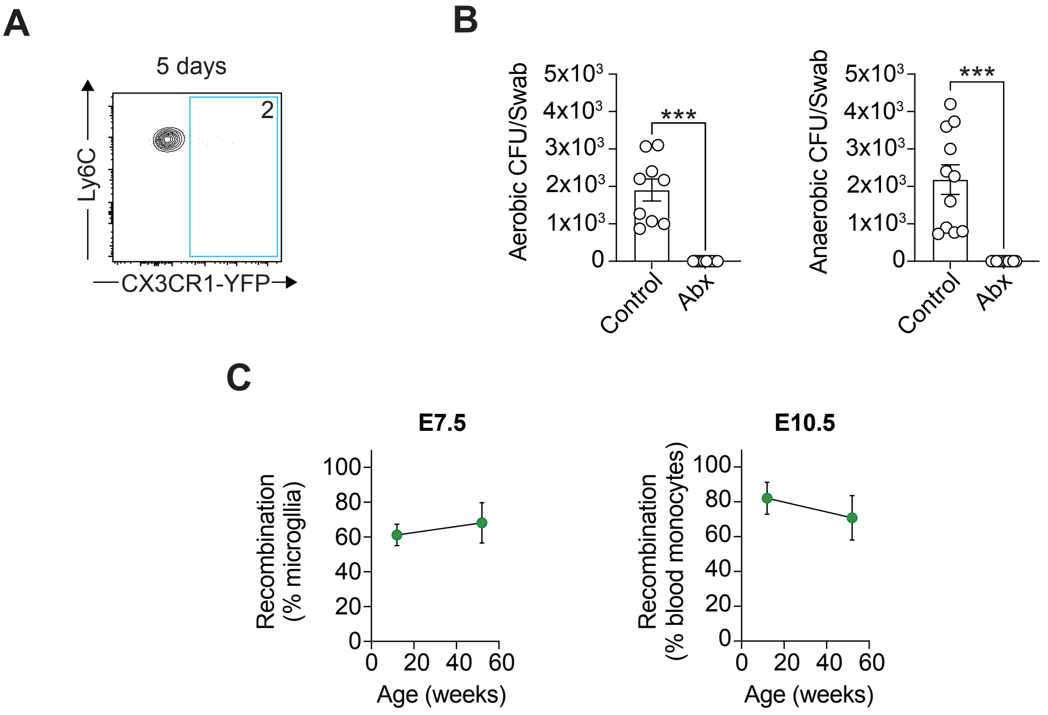


**Figure S2: Transgenic strategies to track gingiva macrophage ontogeny and turnover.** **(A)** Representative flow cytometry plot showing the frequency of YFP-expressing gingiva monocytes in *Cx3cr1*$^{CreER}$*xR26-yfp* mice 5 d after tamoxifen treatment. **(B)** Aerobic (left) and anaerobic (right) colony forming units cultured from oral swabs of control and antibiotic (abx) treated animals. **(C)** Frequencies of reporter recombination in microglia (left) and blood monocytes (right) of *Cdh5*-CreERT2:Rosa$^{tdT}$ mice pulsed with 4OHT at E7.5 and E10.5 respectively. Data are representative of at least two individual experiments, n=3-5 per experiment. Error bars show mean ± SEM.

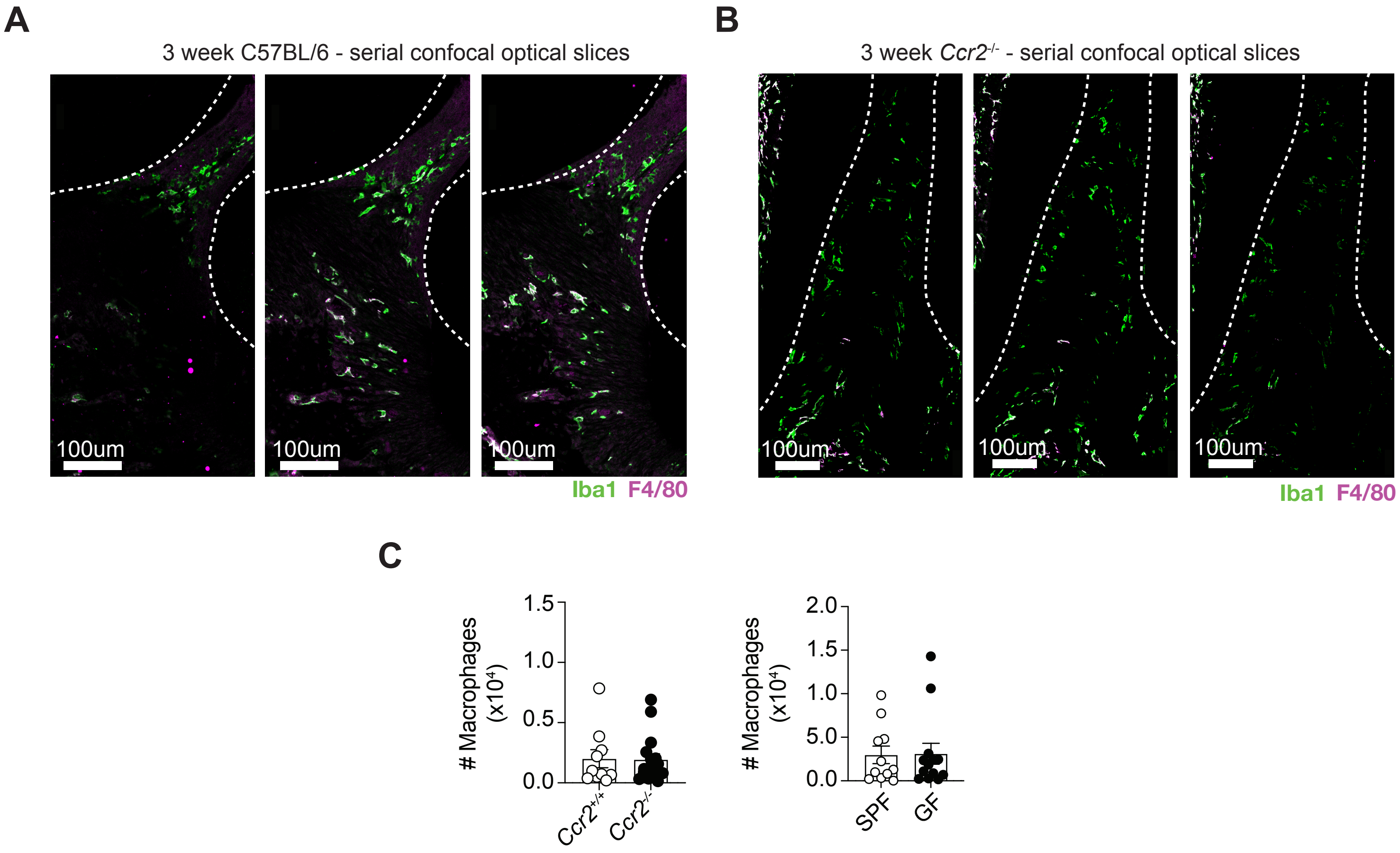


**Figure S3: Adult *Ccr2*$^{-/-}$ and Germ Free mice exhibit unaltered macrophage numbers.**
**(A)** Individual confocal optical sections of Iba1 and F4/80 in gingiva of 3-week old C57BL/6 WT mice. White dotted line indicates outline of tooth. **(B)** Individual confocal optical sections of Iba1 and F4/80 in gingiva of 3-week old *Ccr2*$^{-/-}$ mice. White dotted line indicates outline of tooth. **(C)** Graphs show number of gingiva macrophages in adult *Ccr2*$^{+/+}$ and *Ccr2*$^{-/-}$ mice (left) and SPF and GF mice (right). Data are representative of at least two individual experiments, n=3-5 per experiment. Error bars show mean ± SEM.

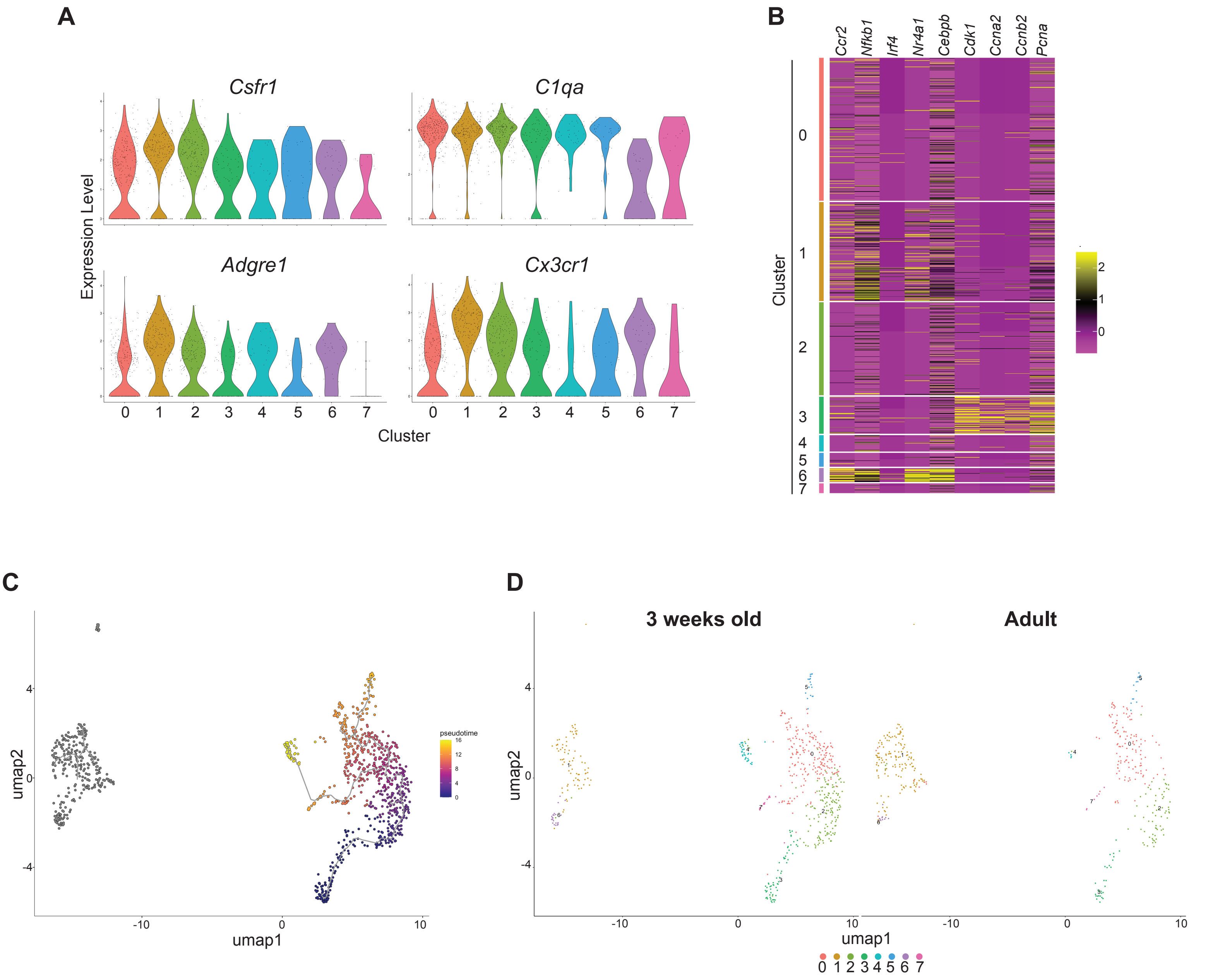


**Figure S4: scRNA-sequencing of gingiva macrophages reveals 8 distinct clusters.**
Gingiva macrophages from 3 week old and adult animals were FACS-sorted and underwent scRNA-sequencing. **(A)** Violin plots showing gene expression of macrophage-defining genes. **(B)** Heat map showing expression of indicated genes in each macrophage cluster. **(C)** RNA velocity pseudotime was projected onto the UMAP shown in Fig. 4A. Dark blue marks the starting point and yellow the end points of the trajectory. **(D)** UMAP of cells from scRNA-seq of sorted macrophages from gingiva of 3 week old (Left) and adult (Right) mice.

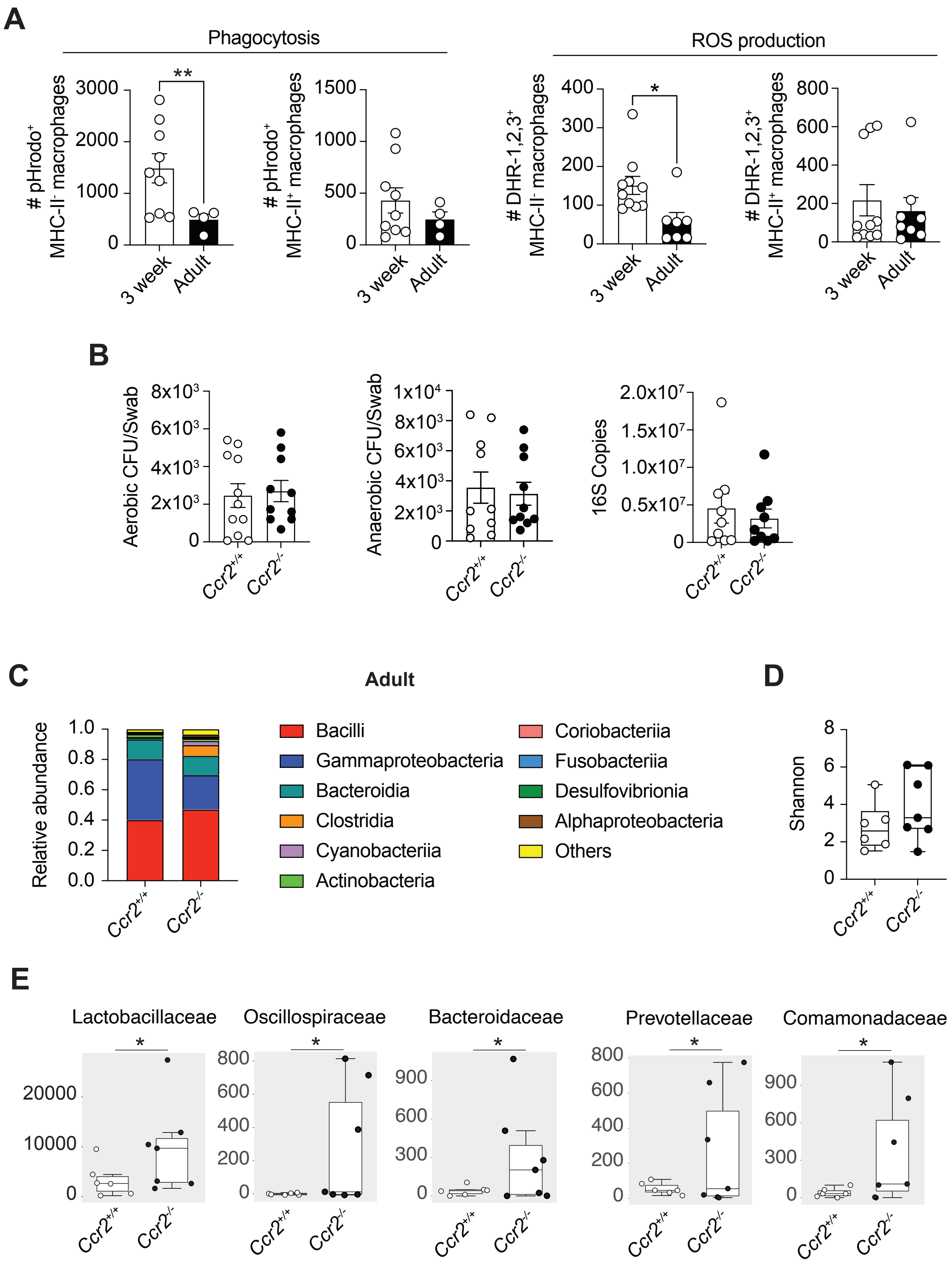


**Figure S5: Functional activities of gingiva macrophages in 3 week old animals**.
**(A)** Total number of DHR-1,2,3$^+$ and pHrodo$^+$ MHC-II$^+$ and MHC-II$^-$ gingival macrophages in 3-wk and 8-10-wk-old C57BL/6 WT mice. **(B)** Aerobic (left) and anaerobic (centre) colony forming units cultured from oral swabs of adult *Ccr2*$^{+/+}$ and *Ccr2*$^{–/–}$ mice. 16S copies as determined by qPCR of DNA isolated from from oral swabs of adult *Ccr2*$^{+/+}$ and *Ccr2*$^{–/–}$ mice (right). **(C)** Relative abundance of bacterial species isolated from oral swabs of adult *Ccr2*$^{+/+}$ and *Ccr2*$^{–/–}$ mice. **(D)** Shannon diversity index of bacterial species isolated from oral swabs of adult *Ccr2*$^{+/+}$ and *Ccr2*$^{–/–}$ mice. **(E)** Relative abundance of bacterial families in adult *Ccr2*$^{+/+}$ and *Ccr2*$^{–/–}$ mice. Data are representative of at least two individual experiments, n=3-5 per experiment. Error bars show mean ± SEM. Statistical comparisons were performed by two-way ANOVA (A): *, P ≤ 0.05, **, P ≤ 0.01.
